# Plastic food packaging from five countries contains endocrine and metabolism disrupting chemicals

**DOI:** 10.1101/2023.09.28.559713

**Authors:** Sarah Stevens, Molly Mcpartland, Zdenka Bartosova, Hanna Sofie Skåland, Johannes Völker, Martin Wagner

## Abstract

Plastics are complex chemical mixtures of polymers and various intentionally and non-intentionally added substances. Despite the well-established links between certain plastic chemicals (bisphenols, phthalates) and adverse health effects, the composition and toxicity of real-world mixtures of plastic chemicals is not well understood. To assess both, we analyzed the chemicals from 36 plastic food contact articles from five countries using nontarget high-resolution mass spectrometry and reporter gene assays for four nuclear receptors that represent key components of the endocrine and metabolic system. We found that chemicals activating the pregnane X receptor (PXR), peroxisome proliferator receptor gamma (PPARγ), estrogen receptor alpha (ERα) and inhibiting the androgen receptor (AR) are prevalent in plastic packaging. We detected up to 9936 chemical features in a single product but found that each product has a rather unique chemical fingerprint. To tackle this chemical complexity, we used stepwise partial least squares regressions and prioritized and tentatively identified the chemical features associated with receptor activity. Our findings demonstrate that most plastic food packaging contains endocrine and metabolism disrupting chemicals. This shows that plastics are a relevant source of exposure to toxic chemicals and further supports the notion that plastic products designed for food contact cannot be considered safe.

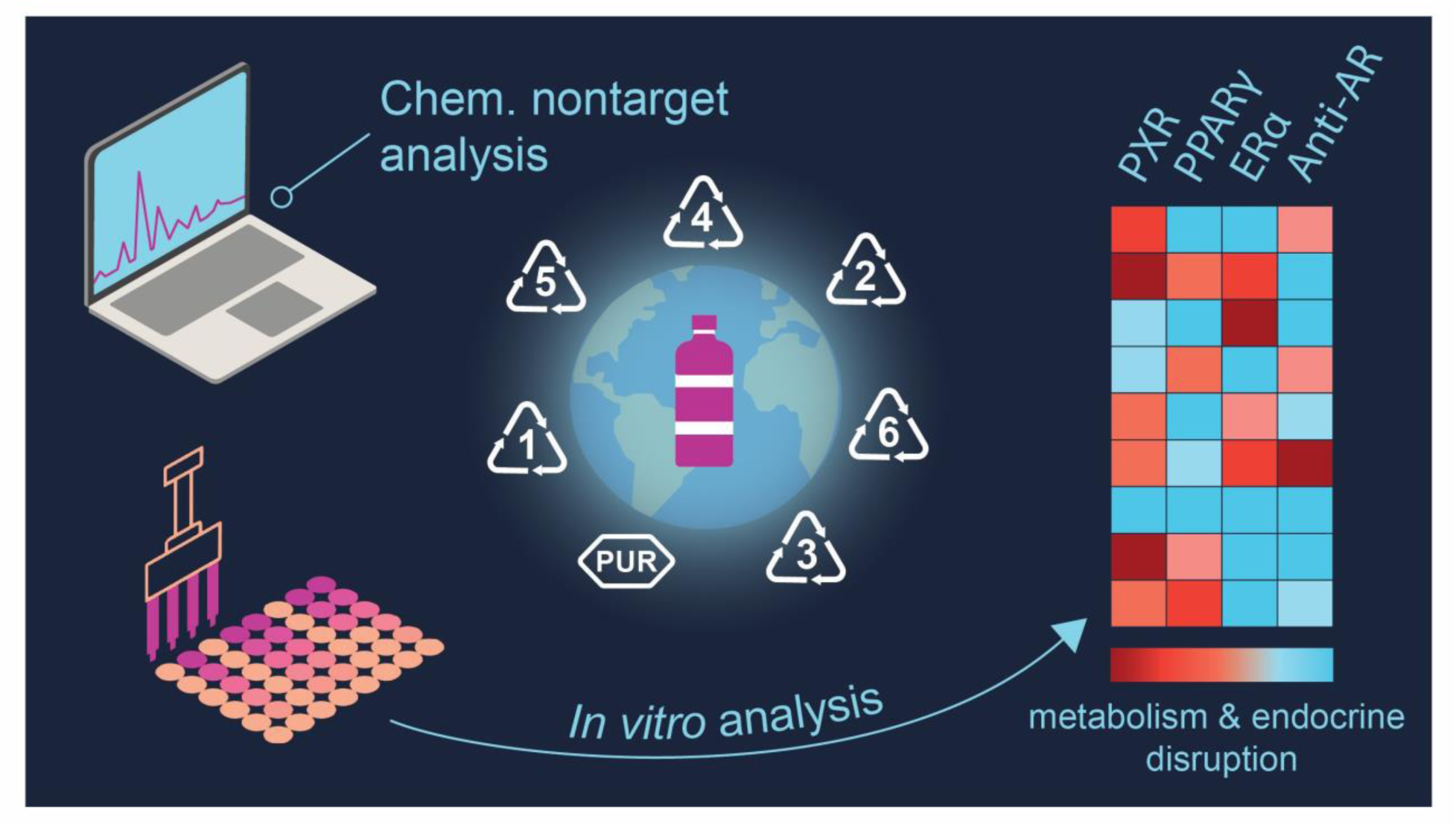

## 1. INTRODUCTION

Plastics are complex mixtures of polymers and a multitude of chemicals used during production either to produce or to enhance the properties of the materials. These chemicals are intentionally added and include residual solvents, monomers, and catalysts as well as a range of additives. In addition, plastic products contain non-intentionally added substances (NIAS), such as impurities and reaction or degradation byproducts generated during production, use, and end of life phase of plastics.^1^ In fact, more than 13 000 plastic chemicals are known.^2–4^

Indeed, plastics are considered a main source of chemical exposures to humans and the environment.^5^ This is because most plastic chemicals are not chemically bound to the polymer matrix, resulting in their release from plastic via migration or volatilization. Within that broader context, plastic food contact articles (FCAs), that is, plastic used to package or process food, is particularly relevant for human exposure.^6^ For instance, certain plastic chemicals, such as bisphenol A (BPA) and phthalates, have been detected in more than 90% of the US population.^7–9^

Plastic chemicals have adverse health effects across the full life cycle of plastics.^10^ Here, endocrine disrupting chemicals, compounds "that interferes with any aspect of hormone action,"^11^ are of particular concern. A chemical disruption of the endocrine system contributes to a wide range of adverse health effects, including reproductive, developmental and metabolic disorders, and cancer.^12^ There is robust evidence that links exposures to BPA and phthalates to such adverse outcomes^13^ resulting in substantial societal costs.^14, 15^ Moreover, emerging evidence suggests that metabolism disrupting chemicals (MDCs) represent another relevant class of compounds.^16, 17^ MDCs promote obesity, type 2 diabetes, or other metabolic disorders, thereby contributing to the increase in non-communicable diseases.^16, 18, 19^ Metabolic disruptions can be mediated via nuclear receptors, such as peroxisome proliferator-activated receptor γ (PPARγ) which is pivotal in lipid metabolism and adipogenesis.^20, 21^ Additionally, pregnane X receptor (PXR), besides its role as xenobiotic sensor, is involved in regulating energy homeostasis, including glucose and lipid and bile acid metabolism.^22–24^ Notably, BPA and phthalates also function as MDCs.^18, 25^ Accordingly, plastic products, including FCAs, can be a source of exposure to EDCs and MDCs.

While much focus is placed on well-studied compounds, the endocrine and metabolism disrupting properties of most plastic chemicals remain unknown. There are multiple reasons for that, including the unresolved identity of many chemicals in plastics, especially of NIAS, and gaps in regulatory frameworks.^26^ Thus, consumers are exposed to mixtures of plastic chemicals with unknown composition and toxicity. Given the vast number of chemicals present in real-world plastic products, bioassays represent a powerful tool to assess the joint toxicity of such complex mixtures^27^ and can be combined with nontarget mass spectrometry.^28^ In our previous work, we applied such an approach and demonstrated that thousands of mostly unknown chemicals are present in a single plastic product which induce a range of toxicological responses *in vitro*.^29^

Nontarget analysis (NTA) provides a comprehensive characterization of the chemical composition of plastics, yet it produces vast amounts of data that are challenging to interpret. This is because high-resolution mass spectrometry generates data on thousands of chemical features in a sample rendering it difficult to pinpoint and identify the active compounds. Statistical models for data reduction, such as partial least squares (PLS) regression, can help address this challenge. Unlike the traditional multiple linear regressions, PLS regression models can manage datasets with numerous co-varying variables and limited sample sizes.^30, 31^ Here, a stepwise method based on variable influence on projection (VIP) scores can be used to select variables that are of importance for each PLS component.^31^ This approach has been successfully implemented by Hug et al. to characterize the chemicals in wastewater effluents inducing mutagenicity.^32^ Thus, integrating bioassays with NTA and multivariate statistics can improve our understanding of chemical mixtures in plastics.

Given our limited knowledge about EDCs and MDCs in plastics, this study aims to investigate the receptor activity induced by all chemicals present in plastic FCAs, representing a real-world scenario that covers unknown compounds and mixture effects. We characterized the receptor activity of all chemicals extracted from plastic FCAs of multiple polymer types in a set of reporter gene assays relevant to human health covering PXR, PPARγ, estrogen receptor α (ERα) and androgen receptor (AR). To gain a more representative picture of the FCAs used globally, we analyzed 36 FCAs from the countries with the highest plastic waste generation per capita.^33^ Further, we used NTA to quantify the chemical features and tentatively identify the chemicals present in the FCAs. We employed PLS regressions as an approach to handle the large chemical complexity encountered in the FCAs and explore a potential relationship between the chemical features and the receptor activity of the samples. Our study confirms the widespread presence of EDCs and MCDs in plastic FCAs and led to the identification and prioritization of several known and unknown chemicals.

## 2. MATERIALS AND METHODS

### 2.1 Samples

We purchased 36 plastic FCAs covering the seven polymer types with the highest global market share,^34^ including high- and low-density polyethylene (HDPE, LDPE), polyethylene terephthalate (PET), polypropylene (PP), polystyrene (PS), polyurethane (PUR), and polyvinyl chloride (PVC) from domestic retailers in five countries (USA, UK, South Korea, Germany, and Norway). We selected four of these countries because of their high plastic consumption, using the plastic waste share per capita as a proxy,^33^ and included Norway because of a local interest. Five to twelve items per country were purchased between winter 2020 and spring 2021. The samples consist of single-use packaging (cups, films, trays, etc.) and FCAs for repeated use (food containers, hydration bladders, etc.). Seventeen FCAs contained food, which was removed by washing with tap water at the local sites. To check for the influence that food content has on chemical composition and toxicity, three samples were acquired, once without and once with food content. All samples were transported to the laboratory in PE bags (VWR) and their polymer type was determined using Fourier-transformed infrared spectroscopy and differential scanning calorimetry in the case of HDPE and LDPE (see S.I. S1.1).

### 2.2 Sample extraction

Methanol (99.8%, Sigma-Aldrich) was used to extract the chemicals from the FCAs as it allows for the extraction of compounds with a large polarity range and does not dissolve the polymers. To avoid sample contamination, consumables used for the extraction were made of glass or stainless steel, rinsed with ultra-pure water (18.2 MΩ cm, PURLAB flex, ELGA) and acetone and heated for at least 2 h at 200 °C. The only exception was the transfer of volumes <1 mL with plastic pipette tips. Prior to extraction, the FCAs which had food content and FCAs of repeated use were rinsed with ultra-pure water and air-dried.

13.5 g of each FCA was cut into smaller pieces (0.5–0.8 x 2 cm, thickness ≤ 0.4 cm) and extracted with 90 mL methanol in two 60 mL glass vials with polytetrafluoroethylene lined lids (DWK Life Sciences). Extraction was performed by sonication (Ultrasonic Cleaner USC-TH, VWR) for 1 h at room temperature. Immediately after, 1 mL extract was removed for the chemical analysis and stored in glass vials at -20 °C. For the bioassays, 60 mL extract was transferred to empty glass vials and evaporated under a gentle stream of nitrogen. When reaching about 0.5 mL, 600 μL dimethyl sulfoxide (DMSO) was added and the evaporation was continued until that volume was reached (i.e., 100-fold concentrated extracts). In parallel, four procedural blanks (PB 1–4) not containing plastics but only methanol underwent the same procedure as the samples to control for a potential contamination during the extraction.

### 2.3 Reporter gene assays

We used CALUX reporter gene assays (BioDetection Systems B.V., Amsterdam) for human PXR, PPARγ, ERα, and AR to analyze the extracts’ receptor activity. The assays were performed as described in Völker et al.^35^ with minor modifications (see S.I. S1.2). On each plate, negative controls (assay medium), vehicle controls (assay medium with 0.05% DMSO), and a concentration series of the reference compounds were included (PXR: nicardipine, PPARγ: rosiglitazone, ERα: 17β-estradiol, AR: flutamide, Table S1 and Figure S2.1). The AR assay was conducted in antagonistic mode with 0.5 μM dihydrotestosterone (CAS 521-18-6, Sigma-Aldrich) as background agonist. Plastic extracts were diluted 500-fold in assay medium and analyzed in five concentrations serially diluted 1:2. Accordingly, the highest analyzed concentration was 1.5 mg plastic well^-1^ (PPARγ, ERα, and AR) or 0.75 mg plastic well^-1^ (PXR), which means that the response observed in the assays was caused by the chemicals extracted from that mass of FCA. Cytotoxic samples were further diluted 1:2 until reaching non-cytotoxic concentrations.

After exposure for 23 h, high-content imaging was used to assess cytotoxicity and normalize the reporter gene response. The cells were stained with NucBlue (Thermo Fisher Scientific) for 30 min, imaged (Cytation 5 Cell Imaging Multimode reader, BioTek) and the nuclei were counted (CellProfiler 4.04). A 20% reduction in nuclei count compared to the pooled negative and solvent controls was used as the cytotoxicity threshold. The receptor activity was subsequently analyzed by measuring luminescence (Cytation 5) of the lysed cells for 1 s following injection of 30 μL illuminate mix containing D-luciferin as substrate. Afterwards, the reaction was quenched in each well with 30 μL 0.1 M NaOH. The extracts were analyzed in a minimum of three independent experiments, each with four technical replicates.

### 2.4 Data analysis reporter gene assays

GraphPad Prism (v10, Graph Pad Software, San Diego, CA) and Microsoft Excel for Windows (v2021-2306) was used for analyzing the bioassay data. Receptor activity was expressed as luminescence normalized to the number of cells well^-1^ and then normalized to the dose-response relationship of the reference compound analyzed on the same plate. Negative and solvent control data were pooled, as they were not statistically different (p > 0.05, Kruskal-Wallis with Dunn’s post hoc test), and the mean was set to 0%. The maximal response to the reference compound (100%) was set to the upper plateau of the dose-response relationship calculated using four-parameter logistic regressions. To derive the effect concentrations of the samples, data were not extrapolated. The limit of detection (LOD) was determined as the average luminesce cell^-1^ of the pooled controls plus 3x the standard deviation. Samples that induced a receptor activity >LOD were considered active.

For data visualization (Figure 3), the effect concentrations were normalized to the lowest (100%) and the highest (0%) analyzed concentration. A Spearman rank correlation matrix was calculated between assay endpoints (normalized EC_20/50_) and feature count. A comparison of samples with and without previous food contact was done using Student’s t-tests. To assess if the FCA’s country of origin or polymer type influence receptor activity, EC_20/50_ values of the different categories were compared using Kruskal-Wallis with Dunn’s post hoc tests. A p < 0.05 was considered statistically significant throughout these analyses.

### 2.5 Chemical analysis

The NTA was performed with an ultra-high performance liquid chromatography system (Acquity I-Class UPLC, Waters) coupled to a high-definition hybrid quadrupole/time-of-flight mass spectrometer Synapt G2-S (Waters). The separation was performed on an Acquity UPLC BEH C18 column (150 × 2.1 mm ID, 1.7 μm, Waters) in a linear gradient with water and methanol as mobile phases, both containing 0.1% formic acid (Table S2). The mass spectrometer was equipped with electron spray ionization source operated in positive mode. Data were acquired over the mass range of 50-1200 Da using data-independent acquisition technique in high resolution (35 000, further details in Table S3). Data treatment and compound identification was performed as described previously^36^ with minor modifications (see S.I. S1.3 and S1.4). The PUR and PVC mass spectra and the PE, PET, PP, and PS spectra were processed separately because their different chemical composition prevented a joint retention time alignment. Features (ions with a unique m/z and retention time) that had an abundance of less than 10-fold the highest abundance across PBs and solvents were excluded from further analysis. Additionally, the abundance of the features was corrected by subtracting the maximum abundance of the respective feature detected in the PBs.

### 2.6 Compound identification, toxicity, and use data

The features remaining after filtering were tentatively identified using the Metascope algorithm in Progenesis QI. The experimental spectra were compared with empirical spectra from MassBank (14 788 unique compounds, release version 2021.03) and spectra predicted *in silico*. For the later, four databases were used as previously described.^36^ In addition, we constructed a fifth database containing the plastic chemicals reported by Wiesinger et al.^4^ as described in S1.3. For the identification, the spectra of each feature in the samples were compared to the spectra in the database with a precursor ion tolerance of 5 ppm and a fragment ion tolerance of 10 ppm. The results of the tentative identification were filtered for hits with a score ≥40 choosing the highest score in case of multiple identifications. Compounds comprising these criteria are referred to as tentatively identified and correspond to the identification level 3.^37^

Publicly available toxicity data of all tentatively identified compounds were retrieved from the ToxCast and Tox21 databases as described previously.^29^ 304 of the tentatively identified compounds had entries in the ToxCast database and we extracted activity concentrations 50 (AC_50_) from 45 assays corresponding to the receptors analyzed here (S.I. S1.4, Table S4).

### 2.7 Partial least squares regression

To reduce the chemical complexity and prioritize chemical features that co-vary with the receptor activity, we conducted PLS regressions with the R package “mdatools”.^38^ Here, we included all features detected in ≥3 samples^30^ and that had an abundance higher than the 25% percentile across all features. The effect concentrations were normalized to the lowest (100%) and the highest (0%) analyzed concentration of the samples. For each receptor, a separate model was applied. The PVC and PUR samples were excluded because three very cytotoxic samples had to be diluted and, thus, could not be analyzed at the same concentrations as the other samples. Through a stepwise variable selection method based on the variables’ influence on projection (VIP), the complexity of the model can be reduced, improving model performance, and selecting features that are important for describing both, the dependent and independent variables.^31^ In an iterative way, we excluded features with a VIP <0.8 until the best model was found or the number of features included in the model was not further reduced.^39^ Model performance was validated by calculating 500 iterations with a random set of features containing the same number of features as in the optimized models. To further identify and prioritize features that positively correlate with the receptor activity, we selected the 25% features which cluster closest to the receptor activity in the first and second component of the optimized model.

## 3. RESULTS AND DISCUSSION

The aim of this study was to assess whether EDCs and MDCs are present in 36 plastic products intended for single or repeated contact with food. We, thus, analyzed whether the chemicals extracted from these products activate or inhibit selected nuclear receptors (PXR, PPARγ, ERα, AR). Additionally, we characterized the chemical composition of the FCAs and prioritized chemical features correlating with receptor activity. To gain a more representative picture of the receptor activity of products consumers use globally, we chose FCAs made of the seven polymers with the highest global production volumes purchased from the countries with the largest plastic waste generation per capita.^33^

### 3.1 Receptor activity of FCA extracts

The chemicals extracted from 33 of 36 plastic FCAs interfered with one or more receptors (Figure 1). We detected PXR agonists in 33 products and PPARγ agonists in 23 products. Cytotoxicity was less prevalent across the samples (S.I. S2.1, Figure S2). This indicates that plastic food packaging contains chemicals that activate the xenobiotic metabolism and interfere with energy homeostasis and metabolic functions. We detected estrogenic and anti-androgenic compounds in 18 and 14 products, respectively. The four procedural blanks were inactive across all assays and experiments, demonstrating that the sample processing did not result in a contamination with chemicals interfering with the receptors investigated here. Taken together, these results imply that most of the plastic FCAs analyzed in this study contain EDCs and MDCs.

**Figure 1.**
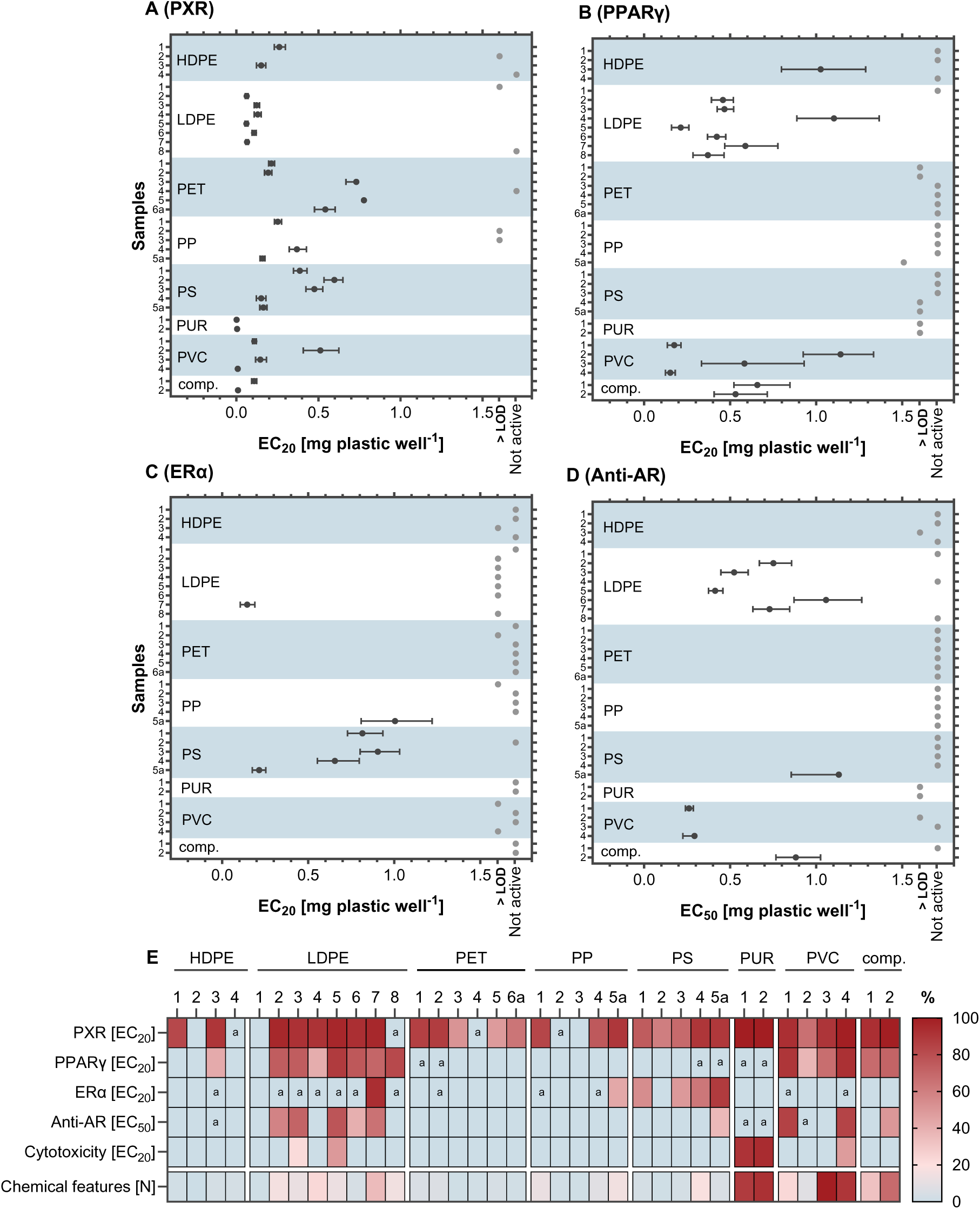
Activity of plastic chemicals extracted from FCAs at (A) PXR, (B) PPARγ, (C) ERα, (D) Anti-AR. (E) Overview of the receptor activity and cytotoxicity normalized to the highest tested concentration, and number of chemical features of the FCAs normalized to the highest detected feature number. EC_20/50_ data are derived from at least three independent experiments, each with four technical replicates per concentration (n ≥ 12). Note: LOD = limit of detection, a = samples with activity between LOD and 20% for PXR, PPARγ and ERα or 50% for Anti-AR.

#### PXR activity

PXR is the predominant target of plastic chemicals extracted from FCAs. All but three extracts activated this receptor and 75% of the extracts produced an EC_20_ (Figure 1A). Three samples, including two PE containers (LDPE 1, HDPE 2) and a PP bowl (PP 3), did not activate PXR nor any other receptor, while eight extracts activated PXR, only. This broad activation of PXR is unsurprising given its promiscuous nature. PXR plays a key role in cellular detoxification and can bind structurally diverse chemicals due to a large ligand binding pocket with several loops in the ligand binding domain.^40, 41^ PXR has important cellular functions beyond its role as xenobiotic sensor, such as energy homeostasis and inflammation.^42–44^ A drug-induced dysregulation of PXR is associated with adverse health effects, including hypercholesterolemia and cardiovascular disease.^45^ Along this line, the plastic chemical dicyclohexyl phthalate was shown to induce PXR-mediated atherosclerosis in mice.^46^ Interestingly, PXR activation correlates positively with the number of chemicals features and with all analyzed biological endpoints, except the estrogenic activity (Figure S3). Screening for PXR activity, therefore, provides a good initial representation of the general toxicity as well as the chemical complexity of mixtures of plastic chemicals.

#### PPARγ activity

The chemicals in 23 FCAs covering every polymer type activated PPARγ (Figure 1B). Compounds extracted from LDPE and PVC products caused a strong receptor activation (>75%), whereas the chemicals in PET, PP and PS FCAs induced effects above the LOD but below 20%. The abundant presence of PPARγ agonists in plastics in this study is interesting since fewer plastic extracts activated that receptor in our previous work.^35^ Thus, PPARγ agonists might be more prevalent in plastic products than previously reported. PPARγ is considered the master regulator of adipogenesis^20^ and its activation by MDCs has been implied in the development of overweight, obesity, and associated metabolic disorders.^18^ Accordingly, we here demonstrate the wide-spread presence of MDCs in plastic FCAs.

#### Estrogenic activity

In total, the chemicals present in 18 FCAs activated ERα (Figure 1C), including four out of five PS samples that induced a potent estrogenic activity (EC_20_ of 0.21–0.91 mg plastic well^-1^). In addition, the compounds in a frozen blueberry package (LDPE 3) and a yogurt cup lid (PP 5a) activated the ERα >20% and another 11 samples induced weak estrogenic effects (>LOD and <20%). This wide-spread detection of estrogenic chemicals in plastics contrasts with our previous work in which 4 out of 34 plastic products contained ERα agonists.^29^ Here, the 10-fold higher sensitivity of the CALUX system compared to the yeast-based reporter gene assay results in lower detection limits and, thus, more detects. However, our findings align with previous studies that observed estrogenic activity leaching from plastic products, including toys.^4748495051^ The prevalence of estrogenic compounds in plastics raises health concerns due to their potential to disrupt the endocrine system, which can, among others, result in developmental and reproductive issues, and an elevated risk of hormone-related cancers, such as breast and prostate cancer.^12^

#### Anti-androgenic activity

We also detected significant anti-androgenicity in 14 samples (Figure 1D). Several LDPE, PVC, and PUR products contained chemicals inducing potent antagonistic effects at the AR, while the compounds extracted from PET and PP articles did not contain anti-androgens. These results are in accordance with Zimmermann et al.^29^ and Klein et al.^52^ who found a similar prevalence of anti-androgenicity in plastics. Further, anti-androgenicity has been detected in plastic baby teethers.^48^ Notably, the anti-androgenic and the PPARγ activities of the extracts correlate significantly (Figure S3) indicating that anti-androgens and metabolism-disrupting chemicals co-occur in FCAs. As for the other nuclear receptors, these results indicate that chemicals with an anti-androgenic mechanism of action are prevalent in plastic food packaging and containers.

Our findings indicate that EDCs and MDCs are frequently present in plastic FCAs from five countries and the compounds present in products of each polymer type interfere with PXR, PPARγ and ERα, and most (HDPE, LDPE, PS, PUR, PVC) inhibited the AR. Nonetheless, we observed some interesting patterns with regards to the polymers: The chemicals in PET and HDPE FCAs activated fewer nuclear receptors (<1/3 active) as compared to the LDPE, PVC, and PUR products (>2/3 active, Figure 1E). Similar to our previous research,^29^ we found individual FCAs made of HDPE, LDPE, PET, and PP that did not contain chemicals interfering with the nuclear receptors. Interestingly, these products also contained very few chemical features (Table 1). This provides two important learnings: (1) it is feasible to produce plastic FCAs from an array of materials that do not contain EDCs or MDCs, and (2) simpler chemical formulations are key to achieving this.

**Table 1.** Plastic food contact articles analyzed in this study and number of chemical features.

| **Polymer** | **Sample name** | **Product** | **Previous food contact<sup>a</sup>** | **Country** | **Chemical features<sup>b</sup>** |
| --- | --- | --- | --- | --- | --- |
| HDPE | HDPE 1 | chewing gum tray | yes | Germany | 164 |
|  | HDPE 2 | food container | no | Germany | 37 |
|  | HDPE 3 | freezer bag | no | South Korea | 260 |
|  | HDPE 4 | milk bottle | yes | USA | 297 |
| LDPE | LDPE 1 | lid food container | no | Germany | 74 |
|  | LDPE 2 | zip lock freezer bag | no | Germany | 1588 |
|  | LDPE 3 | zip lock freezer bag | no | South Korea | 1211 |
|  | LDPE 4 | sausage packaging | yes | South Korea | 2317 |
|  | LDPE 5 | zip lock freezer bag | no | UK | 1243 |
|  | LDPE 6 | cling film | no | USA | 366 |
|  | LDPE 7 | frozen blueberry bag | yes | USA | 3474 |
|  | LDPE 8 | fruit netting | yes | Norway | 1504 |
| PET | PET 1 | oven bag | no | Germany | 528 |
|  | PET 2 | water bottle | yes | South Korea | 640 |
|  | PET3 | dairy cup | yes | UK | 140 |
|  | PET 4 | oven bag | no | UK | 231 |
|  | PET 5 | water bottle | yes | USA | 379 |
|  | PET 6a | food container | no | Norway | 174 |
|  | PET 6b | food container | yes | Norway | 251 |
| PP | PP 1 | dairy cup | yes | Germany | 1220 |
|  | PP 2 | instant food cup | yes | Germany | 194 |
|  | PP 3 | bowl | no | South Korea | 90 |
|  | PP 4 | coffee cup | yes | South Korea | 751 |
|  | PP 5a | yoghurt cup lid | no | USA | 1464 |
|  | PP 5b | yoghurt cup | yes | USA | 1327 |
| PS | PS 1 | dairy cup | yes | Germany | 326 |
|  | PS 2 | bowl | no | Germany | 124 |
|  | PS 3 | cup | no | South Korea | 264 |
|  | PS 4 | cup<sup>c</sup> | no | USA | 1752 |
|  | PS 5a | tray<sup>c</sup> | no | Norway | 504 |
|  | PS 5b | tray<sup>c</sup> | yes | Norway | 1516 |
| PUR | PUR 1 | hydration bladder | no | Germany | 8762 |
|  | PUR 2 | hydration bladder | no | Norway | 9175 |
| PVC | PVC 1 | drinking tube | no | Germany | 2283 |
|  | PVC 2 | drinking tube | no | Germany | 769 |
|  | PVC 3 | cling film | no | UK | 9936 |
|  | PVC 4 | cling film | no | USA | 8946 |
| PET/PE | comp. 1 | sausage packaging | yes | South Korea | 3203 |
| PUR/PE | comp. 2 | cheese packaging | yes | UK | 6803 |

Note: <sup>a</sup> these items had contact with food prior to analysis, <sup>b</sup> number of chemical features detected in nontarget mass spectrometry, <sup>c</sup> items made of extruded polystyrene. Comp. = composite material consisting of two polymers.

### 3.2 Chemical composition of plastic FCAs

#### Individual samples contain a wide range of chemical features

Using high-resolution mass spectrometry, we detected 16 846 unique chemical features in seven PUR and PVC samples and 8665 features in the 29 PE, PET, PP and PS samples. The number of features differed markedly between FCAs with a minimum of 37 features in a food container (HDPE 2) to a maximum of 9936 features present in a cling film (PVC 3, Figure 2A, Table 1). Across all samples, the median number of features was 696 and one quarter of the samples had ≤238 or ≥2150 features, most of which produced robust mass spectrometry signals (Figure 2B). Also, the chemical fingerprints, that is, the presence and abundance of features in each sample, varied greatly between the FCAs (Figure 2C-D). None of the features was detected across all samples in the PE, PET, PP and PS set while 42% (3592 features) are present in only a single FCA (Table S5). Among the samples made of PUR and PVC larger clusters of abundant features are shared between samples (Figure 2D).

**Figure 2.**
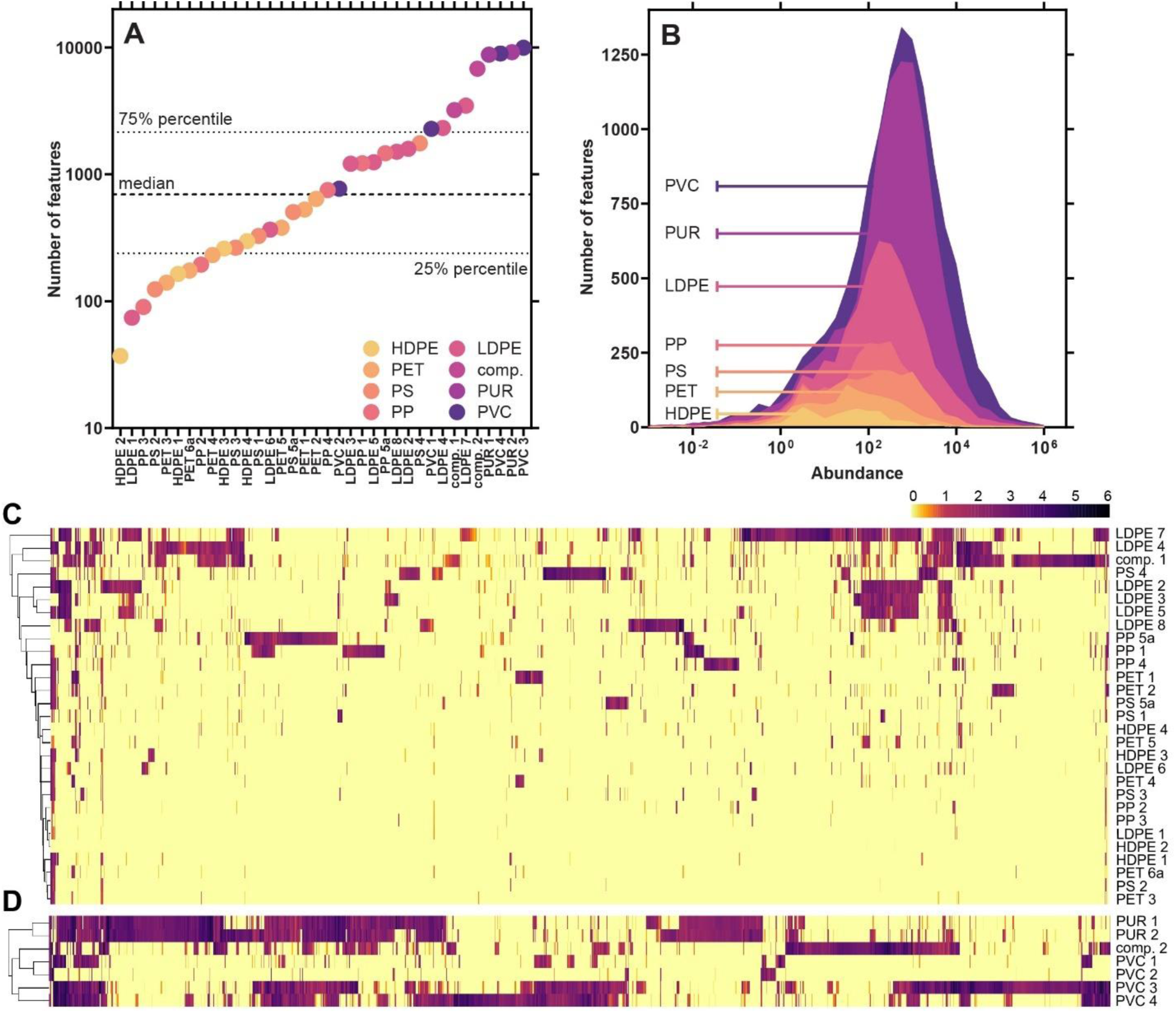
Chemical composition of plastic FCAs. (A) Number of chemical features per sample, (B) abundance of features per polymer type (excluding composite samples), clustered heatmap of chemical features for (C) PE, PET, PP, PS samples and (D) PVC and PUR samples.

**Figure 3.**
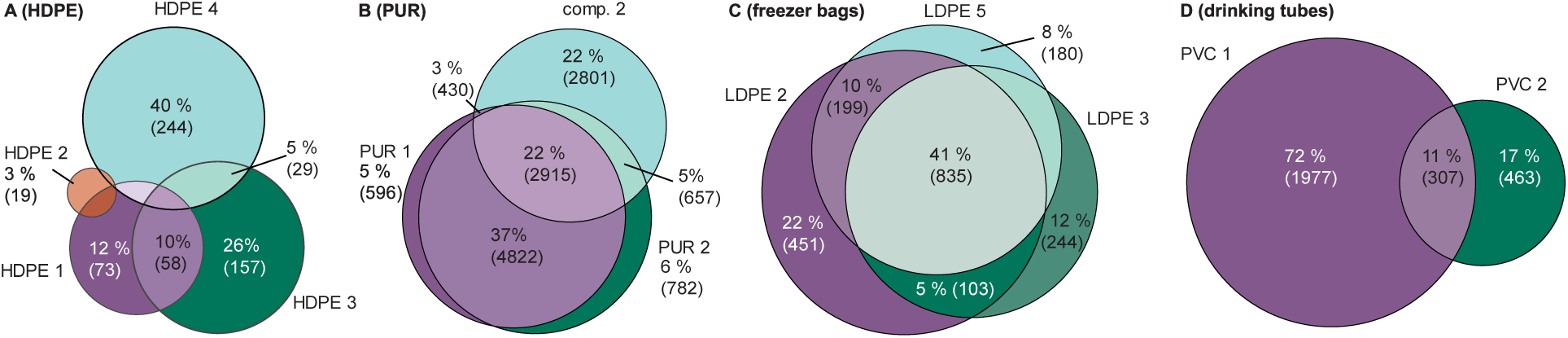
Overlap of chemical features in (A) HDPE samples, (B) PUR samples including the PE-PUR composite (comp. 2), (C) LDPE freezer bags, (D) PVC drinking tubes. An overlap of <1% is not shown.

These results are in line with prior research reporting large numbers of chemicals in consumer plastics using NTA.^36, 52^ Other studies have reported lower numbers of features in plastic products,^53, 54^ that align with our samples that have less features. Notably, the total number of features as well as the full data analysis parameters are rarely reported in nontarget studies of plastic chemicals, somewhat limiting our ability to put our results into context. Nonetheless, the fact that FCAs contain hundreds to thousands of features highlights one dimension of the chemical complexity of plastics, namely the presence of a plethora of chemicals.

#### Polymers differ in number of chemical features

The number of features differs across the polymer types with a gradient ranging from HDPE (616 unique features), PET (1320), PS (2284), PP (2711), LDPE (5495), PVC (12 683) to PUR (13 004, Table S6). We observed a similar pattern with regards to the features’ abundance in the mass spectrometry: the median abundances in HDPE (26) and PET (43) were significantly lower than in PUR and PVC (439 and 508). This indicates that the latter polymers do not only contain more chemicals but also higher levels of those. This is due to the fact that PVC and PUR require more additives in their production compared to other polymers.^5, 55–57^

#### Polymers are chemically diverse

Despite the general trend of more plastic chemicals being abundant in certain polymer types, we observed a striking heterogeneity in the chemical composition of individual FCAs made of the same polymer (Figure 3A and S4). Typically, these products share <2% of the features and, in fact, most features (44–82%) are unique to a single product of a given polymer (Table S6). Only PUR containing samples are an exception to this, they share 22% of features (Figure 3B). With regard to intentionally added substances, this heterogeneity may be explained by the wide range of additives with similar functionality available on the market.^4^ In terms of NIAS, this finding is more surprising because one would expect similar reaction and degradation products to form in products made of the same polymer type. This indicates that very few common chemicals are indeed used or present in a specific plastic type.

When comparing the chemicals fingerprints, the samples did not cluster according to polymer type in the PE, PET, PP and PS samples (Figure 2C). A frozen blueberry packaging (LDPE 7) has the largest number and abundance of features and is most dissimilar to all other samples. Nonetheless, it shares characteristic clusters of features with the three LDPE zip lock freezer bags (LDPE 2, 3, 5) which form a distinct family on its own. These samples share 875 features (43%) and seven of the ten most abundant features (Figure 3C) pointing towards a joint manufacturer. The seven PVC and PUR-based samples, which comprise less diverse articles, cluster better according to polymer type (Figure 2D). Similar to the LDPE freezer bags, samples of the same polymer and product type tend to share larger fractions of chemical features (76% PUR hydration bladders, 53% PVC cling films). However, the two PVC drinking tubes share only 307 chemical features (11%, Figure 3D). Accordingly, neither the polymer type nor the product type is a good predictor for chemical composition. This demonstrates that products with the same functionality can be produced with different numbers and abundance of chemicals.

#### Tentatively identified compounds

In total, we tentatively identified 4137 chemicals (17% of all features). However, this corresponds to 2146 unique identifications only, indicating that multiple features were identified as the same compound (Table S7). In the PE, PET, PP, and PS samples, 1760 chemicals (20%) were identified, comprising 1182 unique chemicals. Among the FCAs made of PVC and PUR, 2377 features (14%) were tentatively identified corresponding to 1371 unique chemicals (Table S7, Table S14, more details in S2.3).

Of the ten most abundant features per sample, we tentatively identified 69 chemicals and retrieved use and toxicity information from PubChem. Our analysis shows that 43 of these chemicals are probably used in plastics as colorants, plasticizers, flame retardants, antioxidants, and processing aids (Table S8). The remaining features either had no use data (n = 12) or are unlikely to be used plastic but as cosmetics, pharmaceuticals, or pesticides (n = 14). Among the plastic chemicals, we detected several known toxic and/or persistent and bioaccumulative compounds with high abundances. One such example is the plasticizer and flame retardant triphenyl phosphate (TPP, CAS 115-86-6) that was detected in both PUR hydration bladders (PUR 1 and 2) with high abundance. TPP is very persistent, very bioaccumulative and toxic to aquatic life.^4^ In addition, the compound interferes with all the receptors analyzed here.^58^ This demonstrates that known hazardous chemicals are used and present in plastic FCAs.

### 3.4 Predictors of receptor activity

We considered the factors of previous food contact, country of origin, polymer type, and presence of known active chemicals as potential predictors of the receptor activity and, in addition, used PLS regressions to identify features potentially contributing to it.

#### Impact of previous food content on receptor activity and chemical composition

We analyzed three samples either with or without previous food content to investigate the impact food storage has on toxicity and chemical composition. The overlap of chemical feature in these paired samples ranged from 28% (PS, 1086 features) to 70% (PP, 422 features), with unique features in both conditions (Figure S5A). This indicates the migration of chemicals from food to the packaging and vice versa. If active, these chemicals will confound the bioassay results. We found that previous food contact increases PXR activity by 39% at the highest concentration for the PET and PS FCAs but decreased it by 12% for the PP cups (Figure S5B-D). However, all FCAs without previous food content had a significant activity on their own. The activity at the more specific receptors PPARγ, ERα, and AR was less affected by the food content with an increase of up to 11, 12 and 18%, respectively, and a decrease of estrogenicity for the PP cups (24%, Figure S5).

We demonstrated that chemicals migrating from food into the FCAs can contribute to the receptor activity, particularly for PXR. This should be considered when testing FCAs, but it can be challenging for researchers to gain access to FCA on the market that were not in contact with food.

However, the previous food content had no significant effect on receptor activity across all extracts (Figure S6). Thus, while food content can be a confounding factor, the general trends observed in receptor activity cannot be attributed to chemicals originating from the food.

#### The country of origin does not influence receptor activity

Across the 36 FCA, the country of origin did not significantly affect the receptor activity (Figure S7). Accordingly, differences in regulations between Germany, Norway, South Korea, the UK, and the US concerning food contact materials or plastic production^59^ do not seem to impact receptor activity. These results are not surprising given the globalized nature of the manufacturing of plastic products.^60^ While the generalizability of our results is limited by the small sample size, these results demonstrate the global dimension of this issue.

#### The polymer type affects the receptor activity

Contrarily to the country of origin, we observed a significant effect of the polymer type on the PPARγ and estrogenic but not on the PXR and anti-androgenic activity (Figure 5, Figure S8). The chemicals in LDPE and PVC induced a significantly stronger PPARγ activity than the ones in PET and PP and the estrogenic activity is significantly stronger in PS than for products made of HDPE, PET, and PVC. This indicates that these estrogenic chemicals are specific to PS products. This is consistent with previous reports on estrogenic compounds in PS migrates^61^ and migrating styrene mono- and oligomers may be causative.^62, 63^ Cytotoxic chemicals are significantly more prevalent in PUR than in the other polymer types, except PVC. This might be due to the large number of chemicals present in both polymers or due to the presence of residual, toxic monomers in PUR.^56, 64^

**Figure 5.**
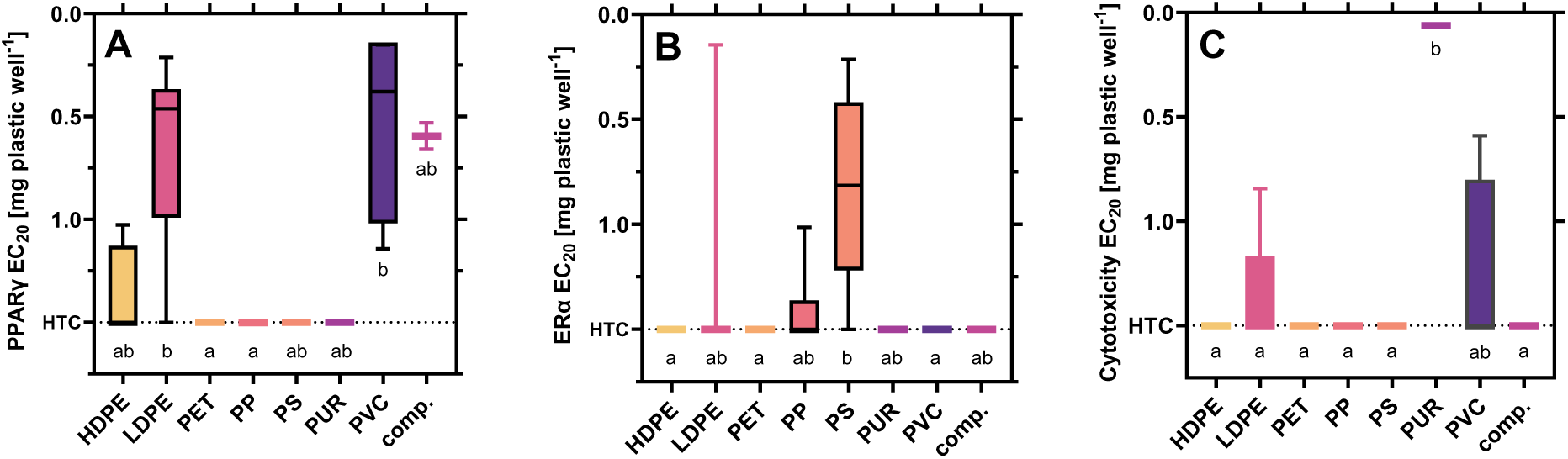
Impact of the polymer type on receptor activity at (A) PPARγ, (B) ERα, and (C) cytotoxicity. EC_20/50_ calculated from at least three independent experiments with four technical replicates per concentration (n ≥ 12). Kruskal-Wallis tests with Dunn’s multiple comparison tests for statistical differences (p < 0.05) indicated by letters.

Our results indicate that the polymer type can predict certain receptor activities and cytotoxicity. While we did not find specific polymers that were free of receptor activity, some polymers (LDPE, PS, PVC, PUR) contain more EDCs and MDCs than others. This means that such polymers could be prioritized for redesign or regulation.

#### Presence of known active compounds in FCAs

To explore if the receptor activity can be explained by known active compounds, we compared the 2146 tentatively identified chemicals with their receptor activity using ToxCast data. Here, 304 compounds detected in our samples are listed in ToxCast of which 298 were active at one or more receptors investigated in this study. These include 117 PXR agonists, 51 PPARγ agonists, 76 ERα agonists and 69 AR antagonists. To prioritize the active compounds, we ranked the compounds based on their abundance in samples as a proxy for concentration and their potency at a receptor according to ToxCast. Within the highest-ranking chemicals, we found plastic-related chemicals, such as triphenyl phosphate (CAS 115-86-6) present in 10 samples of six different polymers, octrizole (CAS 3147-75-9) present in both PUR hydration bladders and tributyl 2-acetyloxypropane1,2,3-tricarboxylate (CAS 77-90-7) present in three samples (LDPE 2, PS 5a, comp. 2, Table S9, further information in S2.5). These results highlight that chemicals with known receptor activity are present in plastic FCAs from across the globe.

To further investigate whether these known active chemicals would predict the observed receptor activity, we compared the detection and the bioassay results. The lack of a clear pattern between the presence of active chemicals in the samples and their respective activity (Table S10) indicates that known active chemicals cannot explain the observed effects. A limitation of this approach is that it is limited to known compounds and ignores their potency and concentration as well as mixture effects. To account for these limitations, we employed PLS regressions, including all chemical features to explore a potential relationship between the occurrence and abundance of chemical features and the receptor activity of the samples.

#### Data reduction to identify relevant chemical features

We used PLS regressions to handle the large chemical heterogeneity of the samples and identify features co-varying with receptor activity. Through a stepwise exclusion of features, we optimized the PLS models for PXR, PPARγ, ERα, and Anti-AR (Figure 6, Table S11). The optimization process reduced the number of features to 1533, 729, 332, and 661 features, respectively, that were identified as important contributors to the receptor activity (Table S12). Compared to the original number of all detected features (8819), this approach reduces the chemical complexity by 82–96%. The optimized models performed better than the initial models and a randomly selected set of features used for validation. The model for PXR activity (Figure 6A and E) resulted in the lowest feature reduction (51%) and the lowest predictive power (cross-validated R^2^ of 0.61) suggesting that multiple compounds contribute to receptor activation in line with the promiscuous binding properties of PXR. The ERα model (Figure 6 C and G) exhibited the best performance, with a pronounced improvement of the cross-validated RMSE and good predictability (cross-validated R^2^ of 0.9), along with a large reduction in the number of features (89% reduction). This indicates that specific chemicals are present in the active samples, significantly co-correlating with their estrogenicity, while these components are absent in the inactive samples. These results align well with the significantly stronger estrogenic activity of PS samples, pointing to the presence of specific estrogenic compounds related to the polymer PS.

**Figure 6.**
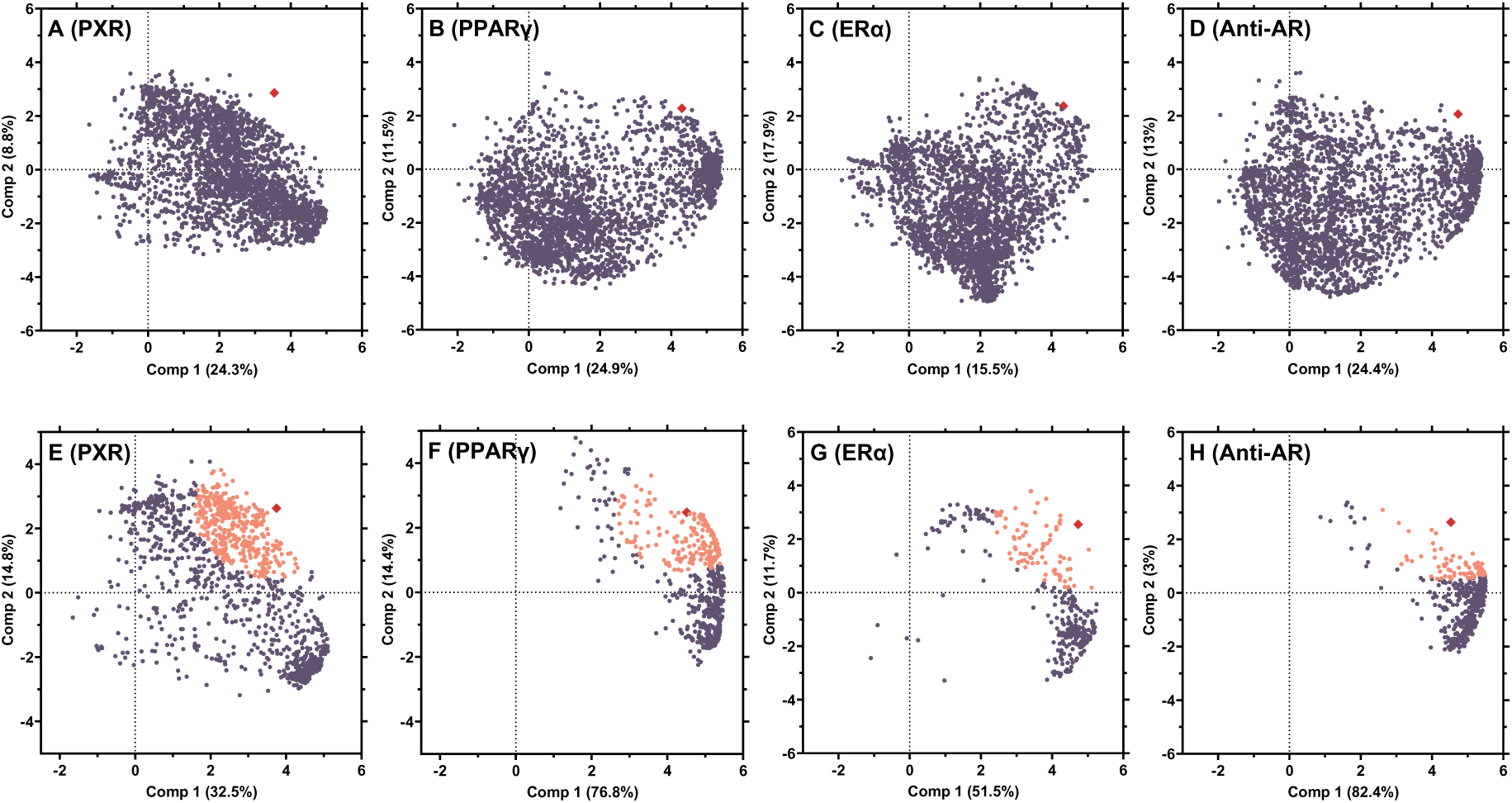
Prioritization of chemical features co-correlating with the receptor activity using stepwise PLS regressions. The analysis refers to food contact articles made of PE, PET, PP and PS. Model 0 (A–D) and the optimized models (E–H) for the respective receptors. The receptor activity is represented by red diamonds, chemical features as circles and features clustering with receptor activity in orange.

To further narrow down to co-varying features that positively correlate with the receptor activity, we selected the 25% features closest to the receptor activity in the optimized model (Figure 6). This additionally reduced the number of features to 383, 95, 83 and 165 for PXR, ERα and AR, respectively. 264 of these features were tentatively identified before, corresponding to 152 compounds, of which 26 are related to plastics^4^ and six were active at the respective receptor according to ToxCast data (Table S13). The latter include the four plastic-related chemicals triethylene glycol (CAS 112-27-6) clustering with PXR, tetradecanoic acid (CAS 544-63-8) with PPARγ, and triphenyl phosphate (CAS 115-86-6) with ERα. The active compound clustering with the anti-androgenic activity, 1-dodecyl-2-pyrrolidinone (CAS 2687-96-9), is a surface-active agent and was detected in all three LDPE freezer bags and the frozen blueberry packaging (LDPE 7).

The co-correlation of the receptor activity and known active compounds strengthens the confidence in using PLS regressions to narrow down the chemical complexity of plastics. In addition, the co-correlation with many other features may contribute to the activity. Here, PLS regressions provide a promising approach to prioritize potentially relevant chemicals that lack identity and activity data for further research.

### 3.5 Implications

Our results confirm that many plastic FCAs contain EDCs and MDCs that interfere with nuclear receptors crucial for human health. The chemicals present in food packaging made of PVC, PUR and LDPE induced most effects whereas the extracts of HDPE, PET, and PP were less active. Nonetheless, we cannot conclude that a particular polymer type is free of toxic chemicals, as samples of each polymer activated most receptors. This research highlights the importance of analyzing the toxicity of whole chemical mixtures of finished plastic products because it covers all extractable chemicals, including unknowns. Using the stepwise PLS regressions, we were able and prioritize chemical features co-varying with the observed receptor activity. This represents an important step towards reducing the chemicals complexity of chemicals plastic products. At the same time, our work also highlights the limited knowledge about the compounds present in plastics. Since many relevant features remain unidentified, we recommend identifying the active compounds in these complex mixtures to enable better monitoring of human exposure and downstream effects. Moving forward, it is essential to consider chemical simplicity as a guiding principle in plastic design and production. This is supported by our findings that chemically less complex plastic products induced a lower toxicity. By using fewer and better characterized chemicals, the safety of plastic products can be significantly improved.

## Acknowledgments

This project has received funding from the European Union’s Horizon 2020 research and innovation program under grant agreement No 860720. Special thanks to Andrea Faltynkova for her valuable help in developing the PLS regression models, to Vaibhav Budhiraja for additional differential scanning calorimetry analyses, and to Dru Jagger, Jaeho Lee, and Mara Mcpartland for their help with obtaining our samples. Your contributions were essential to the success of this study.

## Author contributions

S.S., J.V., and M.W. conceived the study, S.S., M.M., and H.S. preformed the sampling and sample preparation, S.S. performed the bioassay experiments, Z.B. and H.S. performed the chemical analyses, S.S. and H.S. analyzed the data, S.S. and M.W. wrote the manuscript, and all authors provided comments on the manuscript.

## Competing interests

M.W. is an unremunerated member of the Scientific Advisory Board of the Food Packaging Forum Foundation and received travel support for attending annual board meetings.

**Supporting information for this study is available on bioRxiv.**

