## Supporting Information for "Plastic food packaging from five countries contains endocrine and metabolism disrupting chemicals"

**This PDF file includes:**

S1 Supporting methods and materials

S2 Supporting results and discussion

Figures S1 to S8

Tables S1 to S13

Supplementary references

Other supplementary materials for this manuscript include the following:

Table S14-S15

#### S1 Supporting methods and materials

##### S1.1 Determination of polymer type.

The polymer types of the FCA were determined with Fourier-transformed infrared spectroscopy (Cary 610-FTIR and Resolutions Pro software, Agilent) in attenuated total reflectance and transmission mode for the expanded polymers (LDPE 8, PS 5a and b). The polymer type was determined for the inside and outside of the samples. Spectra were acquired with a resolution of 4 $cm^{-1}$ and 32 scans per spectrum and matched to the library of Primpke et al.[1] Where not stated on the packaging, differentiation between HDPE and LDPE products was done by determining the melting point with differential scanning calorimetry (TGA/DSC 1 thermogravimeter, Mettler Toledo) in the temperature range of 25–180 °C. The measurements were carried out in an $N_2$ atmosphere with a 50 mL $min^{-1}$ flow rate and a 10 K $min^{-1}$ heating rate.

##### S1.2 Reporter gene assays

The CALUX cell lines were cultured at 37 °C and 5% $CO_2$ in growth medium consisting of DMEM/F-12 (GlutaMAX, Gibco, 31331093) supplemented with 5% fetal bovine serum (FBS, Sigma Aldrich, F9665), 5.4 mL non-essential amino acids (Sigma Aldrich, M7145), 1% penicillin/streptomycin (Biowest, L0022) and 0.4% geneticin (Sigma Aldrich, A1720). All cell lines were used between passages 3 and 15 and at 70–90% confluency. For the ERα assay, the growth medium was changed to assay medium (DMEM/F-12 with 5% charcoal-stripped FBS (GlutaMAX, Gibco, 11580546), 5.1 mL non-essential amino acids, and 1% penicillin/streptomycin) 2 d prior to the experiments to avoid that the steroids present in normal FBS interfere with the receptor. All assays were conducted with assay medium in 384-well plates (CellStar 781098, Greiner Bio-One). The outer line of the plates (two lines for the Anti-AR assay) contained medium without cells to prevent edge effects on cell growth. 5000 cells $well^{-1}$ were seeded 24–28 h prior to the 23-h exposure.

Table S1. Reference compounds used in the bioassays.

| Receptor | Reference compound | CAS | Supplier | Concentration range [mol L-1] |
|---|---|---|---|---|
| PXR | nicardipine hydrochloride | 54527-84-3 | Sigma-Aldrich | $9.4 \times 10^{-9}$ - $4.8 \times 10^{-6}$ |
| PPARγ | rosiglitazone | 122320-73-4 | Sigma-Aldrich | $3.7 \times 10^{-10}$ - $3 \times 10^{-6}$ |
| ERα | 17-beta-estradiol | 50-28-2 | Sigma-Aldrich | $1.9 \times 10^{-13}$ - $1.0 \times 10^{-10}$ |
| AR | flutamide | 13311-84-7 | Sigma-Aldrich | $1 \times 10^{-8}$ - $3 \times 10^{-5}$ |

#### S1.3 Chemical analysis.

For chromatographic separation a 38-min linear gradient program was used (Table S1) and compounds eluting between 1.5 and 36 min were included in the analysis. The flow rate was set to 0.2 mL $min^{-1}$, the column temperature was maintained at 55°C and the injection volume was 2 µL. After each sample a 7 min wash step with 100% methanol at 0.35 mL $min^{-1}$ was included, followed by a 3 min equilibration to the initial column conditions. The collision energy ramped from 15 to 45 V, and the scan time was 0.3 s (details in Table S3). The analysis included all samples, PBs 1–4, the solvents used during extraction (methanol evaporated to DMSO), and **7** quality controls (pool of all PUR and PVC or PP, PE, PET, and PS samples).

Data treatment and compound identification was performed with the software Progenesis QI (Nonlinear Dynamics, version 3.0). The retention times were automatically aligned to one of the quality controls (pool of all samples) selected by the software. Peak picking was performed with automatic sensitivity to detect "fewer" peaks. The minimum chromatographic peak width was set to 0.05 min and the fragment sensitivity was set to 0.2% of the base peak. The program searched for common adducts (M+H, 2M+H, M+2H, M+H-$H_2O$, M+Na, and 2M+Na), and deconvoluted the mass spectra. The resulting list of chemical features was exported to Microsoft Excel for Windows (Version 2021-2306).

The R package "pheatmap" [2] was used for the hierarchical cluster analysis (Euclidian distances) of the chemical features (Figure 2C and 4I). Euler diagrams were produced with the R package "eulerr" (Figure 4 and S5).[3]

Table S2. Chromatographic parameters for the nontarget chemicals analysis.

| Retention time (min) | Mobile phase A (%, water with 0.1% formic acid) | Mobile phase B (%, methanol with 0.1% formic acid) |
|---|---|---|
| 0 | 80 | 20 |
| 0.5 | 80 | 20 |
| 29 | 5 | 95 |
| 36 | 5 | 95 |
| 36.1 | 0 | 100 |
| 38 | 0 | 100 |

Table S3. Mass spectrometer parameters for the nontarget chemicals analysis.

| Parameter | Value |
|---|---|
| Capillary (kV) | 2.8 |
| Source temperature (°C) | 120 |
| Sampling cone voltage (V) | 30 |
| Source offset voltage (V) | 50 |
| Desolvation temperature (°C) | 500 |
| Cone gas flow (L $h^{-1}$) | 500 |
| Desolvation gas flow (L $h^{-1}$) | 900 |
| Nebulizer gas flow (bar) | 6 |

#### S1.4 Compound identification, ToxCast and PubChem search

Identification was done with the Metascope algorithm in Progenesis QI. For the *in silico* fragmentation, four databases were used as previously described.[4] In addition, we constructed a fifth database containing the plastic chemicals described by Wiesinger et al.[5] For their PlasticMap database, we converted the available 10 547 CASRNs to SMILES-codes (n = 6631) using the EPA CompTox batch search.[6] The SMILES codes were then converted to CIDs (n = 6423) using the PubChem Identifier Exchange service [7]. CIDs were uploaded to PubChem to retrieve the chemical structures (n = 6414) as a sdf file which was subsequently used for the *in silico* fragmentation search.

Among the tentatively identified compounds 304 were present in the ToxCast data based on matching InChlKeys, IUPAC names or common names. We extracted their activity concentrations 50 ($AC_{50}$) in 45 ToxCast assays that matched the receptors analyzed here (Table S4). ToxCast assays were selected based on the receptor gene names. The $AC_{50}$ values were compared to the compound's highest peak area and ranked based on the $AC_{50}$/area ratio, for all compounds with a peak area >100 (Table S9). A Spearman rank correlation between number of compounds activating a certain receptor and the samples effect concentrations was calculated with GraphPad Prism (9-10).

For the tentatively identified compounds among the ten features with the highest abundance per sample, an additional search for toxicity and usage information was done on PubChem,[8] and in the supplementary information from Wiesinger et. al. and Groh et al.[5,9]

Table S4. Selected ToxCast assays used to retrieve information about PXR, PPARγ, estrogenic and anti-androgenic activity.

| Receptor | Assay name in ToxCast | Mode |
|---|---|---|
| **PXR** | ATG_PXRE_CIS_up | agonist |
| | ATG_PXR_TRANS_up | agonist |
| | NVS_NR_hPXR | agonist |
| | TOX21_PXR_Agonist | agonist |
| | ATG_PXRE_CIS_dn | antagonist |
| | ATG_PXR_TRANS_dn | antagonist |
| **PPARγ** | ATG_PPRE_CIS_up | agonist |
| | ATG_PPARg_TRANS_up | agonist |
| | NVS_NR_hPPARg | agonist |
| | OT_PPARg_PPARgSRC1_0480 | agonist |
| | OT_PPARg_PPARgSRC1_1440 | agonist |
| | TOX21_PPARg_BLA_Agonist_ratio | agonist |
| | TOX21_PPARg_BLA_antagonist_ratio | antagonist |
| | ATG_PPARg_TRANS_dn | antagonist |
| | ATG_PPRE_CIS_dn | antagonist |
| **ERα** | ACEA_ER_80hr | agonist |
| | ATG_ERE_CIS_up | agonist |
| | ATG_ERa_TRANS_up | agonist |
| | NVS_NR_hER | agonist |

| Receptor | Assay name in ToxCast | Mode |
|---|---|---|
| **ERα** | OT_ER_ERaERa_0480 | agonist |
| | OT_ER_ERaERa_1440 | agonist |
| | OT_ERa_EREGFP_0120 | agonist |
| | OT_ERa_EREGFP_0480 | agonist |
| | TOX21_ERa_BLA_Agonist_ratio | agonist |
| | TOX21_ERa_LUC_VM7_Agonist | agonist |
| | TOX21_ERa_LUC_VM7_Agonist_10nM_ICI182780 | agonist |
| | ATG_ERa_TRANS_dn | antagonist |
| | ATG_ERE_CIS_dn | antagonist |
| | TOX21_ERa_BLA_Antagonist_ratio | antagonist |
| | TOX21_ERa_LUC_VM7_Antagonist_0.5nM_E2 | antagonist |
| | TOX21_ERa_LUC_VM7_Antagonist_0.1nM_E2 | antagonist |
| **AR** | ATG_AR_TRANS_dn | antagonist |
| | NVS_NR_hAR | antagonist |
| | OT_AR_ARELUC_AG_1440 | antagonist |
| | OT_AR_ARSRC1_0480 | antagonist |
| | OT_AR_ARSRC1_0960 | antagonist |
| | TOX21_AR_BLA_Antagonist_ratio | antagonist |
| | TOX21_AR_LUC_MDAKB2_Antagonist_10nM_R1881 | antagonist |
| | TOX21_AR_LUC_MDAKB2_Antagonist_0.5nM_R1881 | antagonist |
| | ACEA_AR_antagonist_80hr | antagonist |
| | ATG_AR_TRANS_up | agonist |
| | TOX21_AR_BLA_Agonist_ratio | agonist |
| | TOX21_AR_LUC_MDAKB2_Agonist | agonist |
| | TOX21_AR_LUC_MDAKB2_Agonist_3uM_Nilutamide | agonist |
| | ACEA_AR_agonist_80hr | agonist |

### S2 Supporting results and discussion

#### S2.1 Receptor activity and cytotoxicity

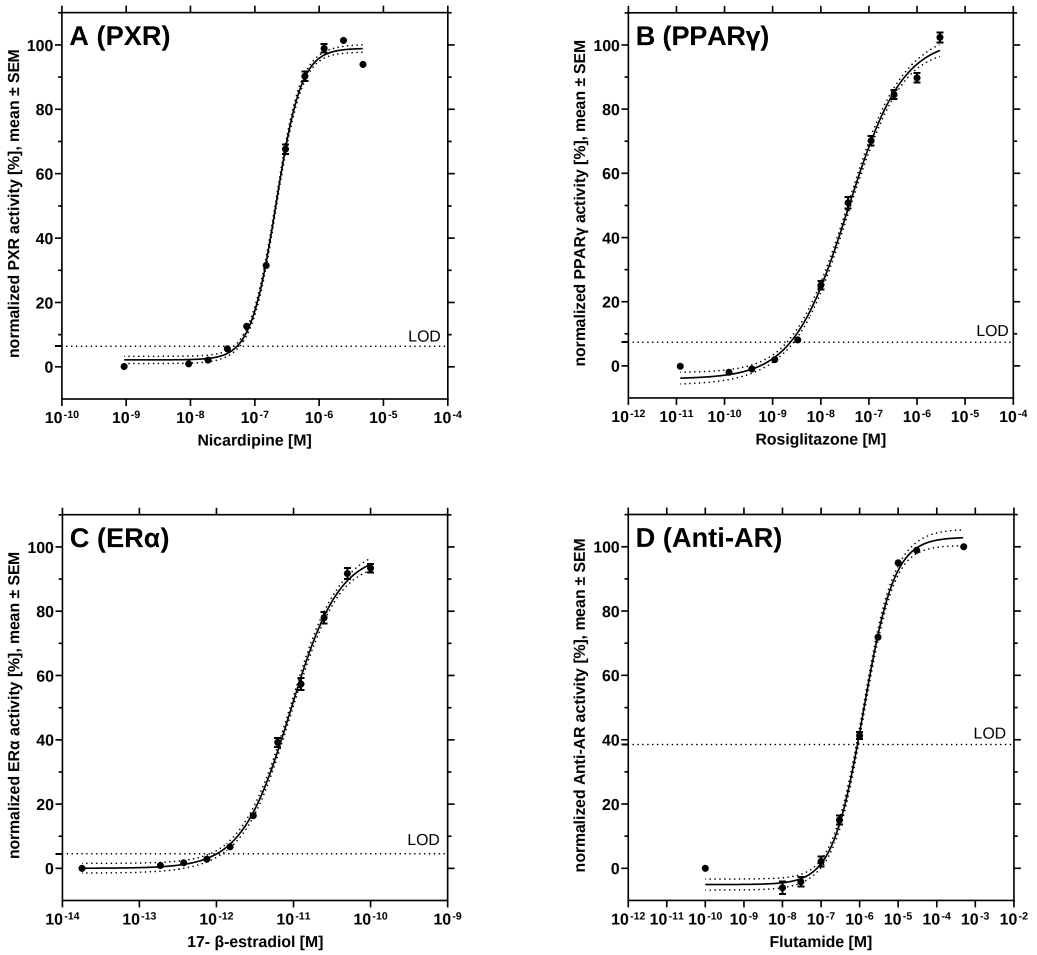


Figure S1. Dose-response relationships of the reference compounds used in the reporter gene assays. Nicardipine was used for the pregnane X receptor (PXR, A), rosiglitazone for peroxisome proliferator receptor gamma (PPARγ, B), 17-β estradiol for estrogen receptor alpha (ERα, C), and flutamide for the anti-androgenic activity (Anti-AR, D). n ≥ 60. Note: LOD = limit of detection

Cytotoxic chemicals were least prevalent in our samples (Figure S2): Five of the FCA extracts induced significant cytotoxicity, including two freezer bags (LDPE 3 and 5), two hydration bladders (PUR 1 and 2), and a cling film (PVC 4). The two PUR hydration bladders induced the strongest cytotoxicity ($EC_{20}$ 0.05 and 0.06 mg/well). Cytotoxicity correlates significantly with the number of chemical features (Figure S1) but this correlation is mostly driven by the two PUR samples. Nonetheless, cytotoxicity is mostly thought of as a result of non-specific effects that will accumulate when more chemicals are present in a sample.[10]

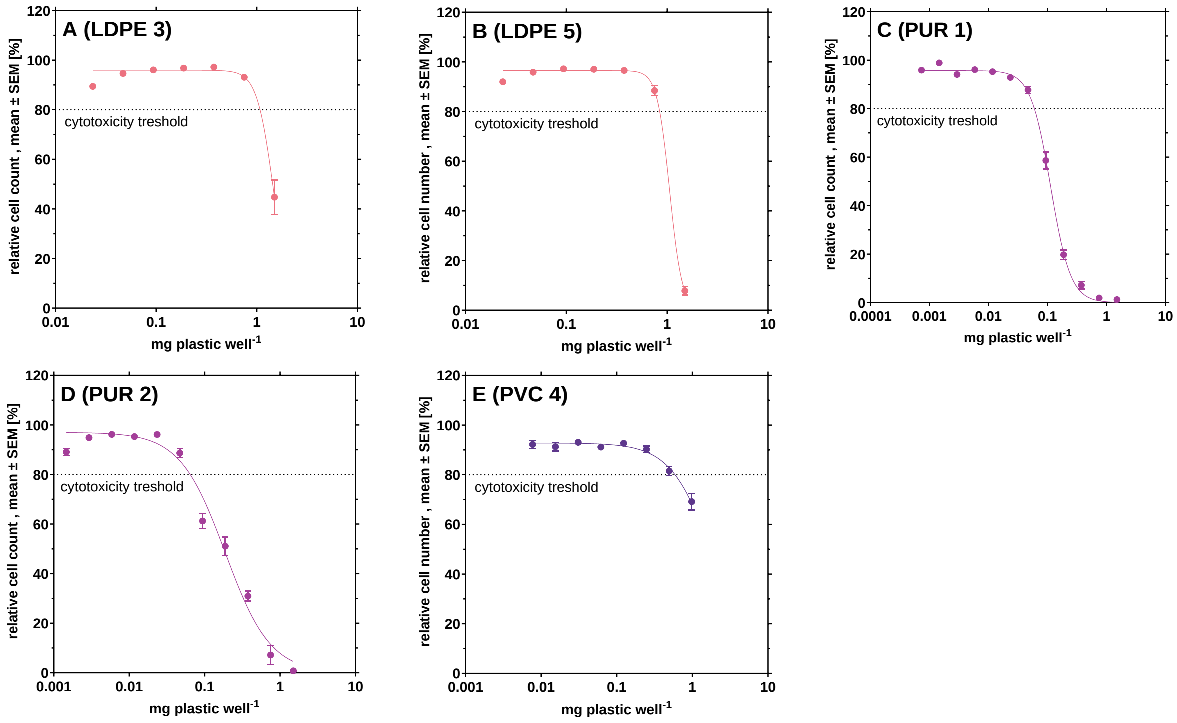


Figure S 2**.** Cytotoxicity of samples LDPE 3 (A), LDPE 5 (B), PUR 1 (C), PUR 2 (D), PVC (4). Cytotoxicity is defined as a cell count <80% of the pooled negative and solvent controls. Cell count across receptors combined, for cytotoxic concentrations n ≥ 16, for non-cytotoxic concentrations n ≥ 48. Highest tested concentration of 1.5 mg plastic/well.

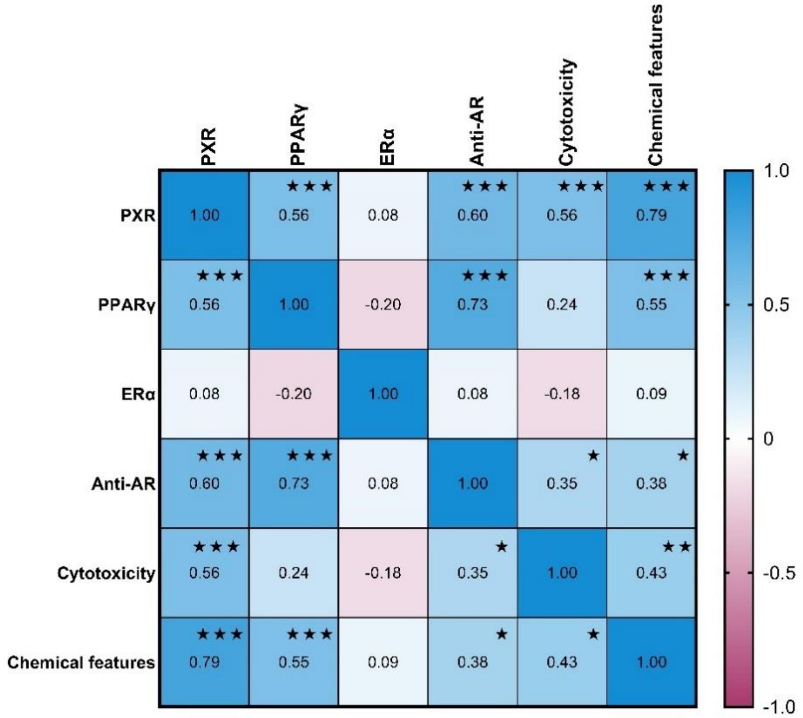


Figure S3. Correlation between biological effects ($EC_{20/50}$) and the number of chemical features detected in the plastic food contact articles. The values in the boxes represent Spearman's rank correlation coefficient, the asterisks indicate that the correlation was statistically significant (* $p < 0.05$, ** $p < 0.01$. *** $p < 0.001$).

#### S2.2 Chemical analysis.

Table S5. Distribution of chemical features across samples.

| PE, PET, PP and PS samples | | | PUR and PVC samples | | |
|---|---|---|---|---|---|
| Number of samples | Number of features shared between samples | Share of all 8819 features [%] | Sample number | Number of features shared between samples | Share of all 16 847 features [%] |
| 1 | 3592 | 41.5 | 1 | 3659 | 21.7 |
| 2 | 1864 | 21.5 | 2 | 4827 | 28.7 |
| 3 | 985 | 11.4 | 3 | 3269 | 19.4 |
| 4 | 773 | 8.9 | 4 | 2583 | 15.3 |
| 5 | 416 | 4.8 | 5 | 1901 | 11.3 |
| 6 | 338 | 3.9 | 6 | 532 | 3.2 |
| 7 | 202 | 2.3 | 7 (all) | 76 | 0.5 |
| 8 | 147 | 1.7 | | | |
| 9 | 88 | 1.0 | | | |
| 10 | 61 | 0.7 | | | |
| 11 | 28 | 0.3 | | | |
| 12 | 33 | 0.4 | | | |
| 13 | 29 | 0.3 | | | |
| 14 | 20 | 0.2 | | | |
| 15 | 11 | 0.1 | | | |
| 16 | 10 | 0.1 | | | |
| 17 | 12 | 0.1 | | | |
| 18 | 16 | 0.2 | | | |
| 19 | 5 | 0.1 | | | |
| > 20 | 35 | 0.4 | | | |
| 29 (all) | 0 | 0.0 | | | |

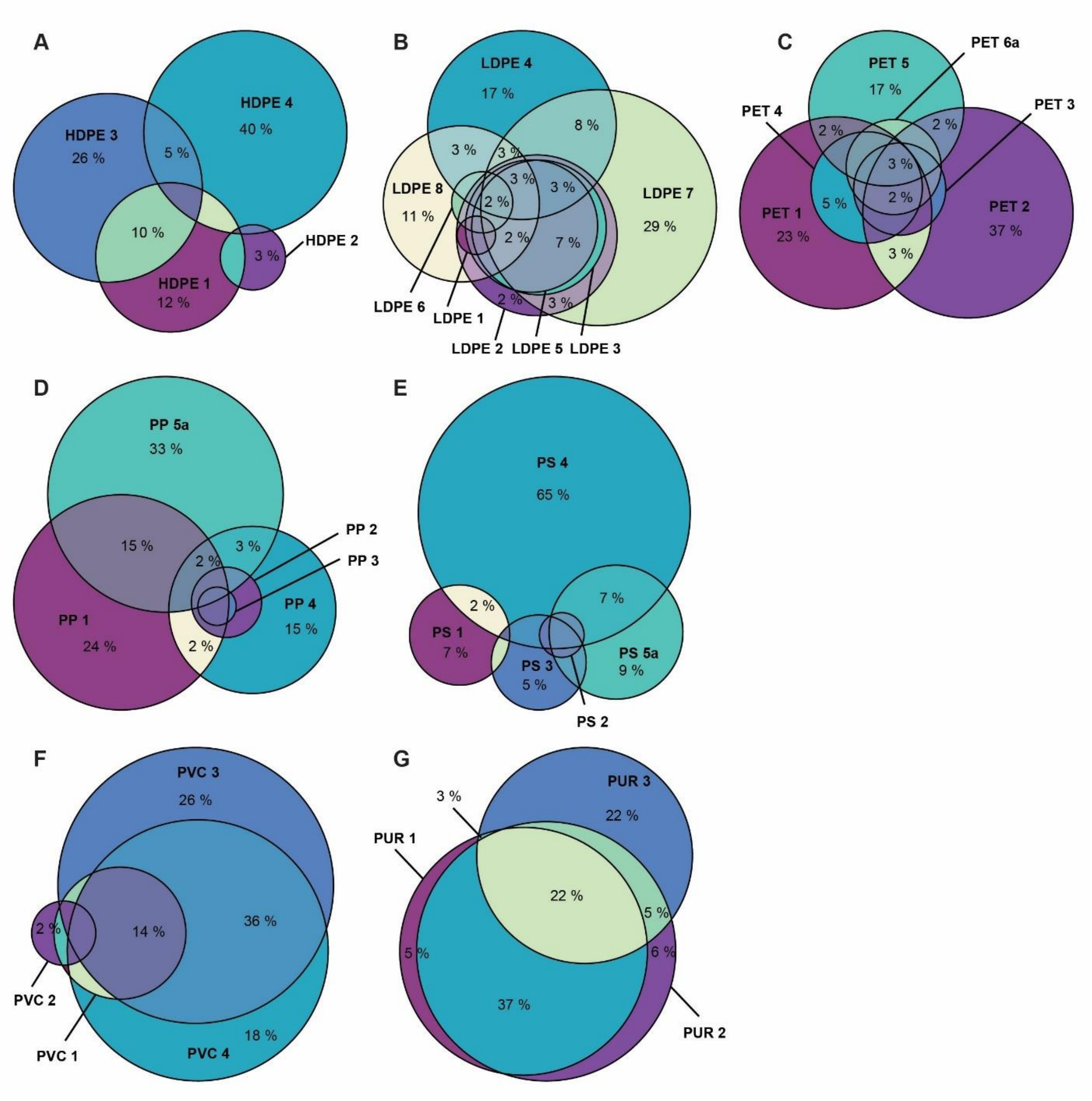


Figure S4. Overlap of chemical features in the samples made of the same polymer (A) HDPE, (B) LDPE, (C) PET, (D) PP, (E) PS, (F) PVC, (G) PUR and PUR-PE composite. An overlap of <1% is not shown.

Table S6. Chemical features in the FCAs made of the same polymer, total number, common features and feature unique to a single product.

| Polymer | N samples | All chemical features | Common chemical features | Unique chemical features |
|---|---|---|---|---|
| **HDPE** | 4 | 616 | 3 (0.49%) | 493 (80.0%) |
| **LDPE** | 8 | 5494 | 2 (0.04%) | 2817 (51.3%) |
| **PET** | 6 | 1320 | 37 (2.8%) | 965 (73.1%) |
| **PP** | 5 | 2711 | 2 (0.07%) | 1929 (71.1%) |
| **PS** | 5 | 2284 | 41 (1.8%) | 1864 (81.6%) |
| **PUR** [a] | 3 | 13 004 | 2915 (22%) | 4180 (32.1%) |
| **PVC** | 4 | 12 683 | 179 (1.4%) | 5738 (44.1%) |

Note: [a] including the composite made of PUR and PE.

##### S2.3 Tentatively identified chemicals

Most of the chemicals were identified with the NORMAN Suspect List Exchange database (77% in the PE, PET, PP and PS samples and 91% in the PUR and PVC samples),[11] which is the most comprehensive database used in this study and includes about 70 000 compounds but is not specific to plastics.

Of the ten most abundant features per sample, we tentatively identified 69 compounds and retrieved use and toxicity information from Pubchem. Our analysis showed that 31 of these compounds were associated with plastics, including colorants, plasticizers, flame retardants, antioxidants, and other processing aids (Table S8). Among the remaining 38 features, 13 have no relevant use or toxicity data while 25 are not related to plastic and consist of cosmetics, pharmaceuticals, and fungicides among others (Table S8). Examples of plastic additives detected are erucamide (CAS 112-84-5) and octadecanamide (CAS124-26-5), which are used as colorants, fillers, and lubricants.[5] These substances were found with high abundance in the three LDPE freezer bags (LDPE 2, 3 and 5). Further, we detected the NIAS ethylene terephthalate cyclic trimer (CAS 7441-32-9) with high abundance in five of the six PET samples. This compound is known to occur in PET.[9]

Among the plastic additives, we detected several known toxic and/or persistent and bioaccumulative compounds with high abundances, such as triphenyl phosphate (TPP, CAS 115-

86-6), octrizole (CAS 3147-75-9), bis(1,2,2,6,6-pentamethylpiperidin-4-yl)decanedioate (CAS 41556-26-7), methyl 1,2,2,6,6-penta methyl- 4-piperidyl sebacate (CAS 82919-37-7) and ethyl 4-[(N-methylanilino)methylideneamino]benzoate (CAS 57834-33-0). The plasticizer and flame retardant TPP has bioaccumulative, persistent and endocrine disrupting properties[8] and interferes with all the receptors analyzed in this study.[12] TPP was detected in both PUR hydration bladders (PUR 1 and 2), while the other four compounds were detected in only one hydration bladder (PUR 2). Tributyl 2-acetyloxypropane-1,2,3-tricarboxylate (CAS 77-90-7) is another toxic plasticizer that was detected in an LDPE and PS sample (LDPE 2 and PS 5b) and is persistent, bioaccumulative, carcinogenic, and mutagenic. [8]

Table S7. Number of features and identified compounds detected in the samples.

| Polymer set | Total number of chemical features | Tentatively identified chemical features | Tentatively identified compounds |
|---|---|---|---|
| PE, PET, PP, PS | 8665 | 1760 | 1182 |
| PUR, PVC | 16 847 | 2377 | 1371 |
| Total | | | 2146 |

Table S8. Tentatively identified chemicals among the top ten features per sample with use and toxicity data retrieved from Groh et al. (2019) comprising plastic chemicals specific to food contact articles (A), Wiesinger et al. (2021) comprising intentionally added plastic chemicals (B), PubChem (C) and ToxCast (D).

| CAS | Name | High abundance in sample | Plastic related | Use | Toxicity |
|---|---|---|---|---|---|
| 23470-00-0 | 2,3-dihydroxypropyl hexadecanoate | LDPE_8 | yes (B) | Cosmetics (C) | Toxic to aquatic life (C) |
| 85391-47-5 | 2-O-(2-ethylhexyl) 1-O-undecyl benzene-1,2-dicarboxylate | HDPE 1, LDPE 4, PET 6b | tentative | Adhesive (C) | - |
| 63753-10-6 | 2-hydroxy-5-nonylbenzaldehyde | HDPE 3 | - | Used in metal extraction process (C) | corrosive, irritant, health hazard (C) |
| 77-90-7 | tributyl 2-acetyloxypropane-1,2,3-tricarboxylate | LDPE 2, PS 5b | yes (B) | Plasticizer, other processing aids, FCA (A, B) | Persistent, bioaccumulate, CMR, carcinogenic, mutagenic (B) PPARγ and PXR agonist (D) |
| 112-84-5 | erucamide | LDPE 2, LDPE 3, LDPE 5 | yes (B) | Colorant, filler, lubricant, FCM (B) | Irritant (C). Not classified as of potential concern (B) |
| 10436-08-5 | icos-11-enamide | LDPE 2 | yes (B) | FCM (B) | Not classified as of potential concern (B) |
| 124-26-5 | octadecanamide | LDPE 2, LDPE 3, LDPE 5, PS 1 | yes (B) | Antistatic Agent, Colorant, Filler, Intermediates, Lubricant, FCM (B) | Not classified as of potential concern (B) |
| 59227-89-3 | laurocapram | LDPE 3 | - | Skin conditioning (C) | Irritant (C). AR antagonist, PPARγ, ERα and PXR agonist (D) |
| 94713-19-6 | 4-(4,7-dimethyl-2,3,3a,4,5,6,7,7a-octahydro-1H-inden-5-yl)-2-methylcyclohexan-1-ol | LDPE 3 | - | - | - |
| 5834-63-9 | 1,8,15,22-tetrazacyclooctacosane-2,9,16,23-tetrone | LDPE 4 | yes (A, B) | - | - |
| 621-70-5 | 2,3-di(hexanoyloxy)propyl hexanoate | LDPE 4, comp. 1 | tentative | Lubricants and lubricant additives, textile manufacturing (C) | - |

| CAS | Name | High abundance in sample | Plastic related | Use | Toxicity |
|---|---|---|---|---|---|
| 30899-62-8 | 2,3-diacetyloxypropyl dodecanoate | LDPE 4, comp. 1 | yes (B) | plasticizer, biocide, colorant, filler, lubricant, other processing aids, cling films (B, C) | Not classified as of potential concern (B) |
| 33599-07-4 | 2,3-diacetyloxypropyl octadecanoate | LDPE 4, comp. 1 | yes (B) | Plasticizer in PVC (B) | Not classified as of potential concern (B) |
| 56149-06-5 | 2,3-diacetyloxypropyl docosanoate | LDPE 4 | - | - | - |
| 537-40-6 | 2,3-bis[[(9Z,12Z)-octadeca-9,12-dienoyl]oxy]propyl (9Z,12Z)-octadeca-9,12-dienoate | LDPE 5 | - | solvent, cosmetics (C) | - |
| 25401-86-9 | 2-hexadecylphenol | LDPE 5 | yes (C) | polycarbonate resins (C) | - |
| 94291-86-8 | 8-(3,7-dimethyloct-6-enoxy)-8-methoxy-2,6-dimethyloctan-2-ol | LDPE 6 | - | - | - |
| 112-99-2 | N-octadecyloctadecan-1-amine | LDPE 7, comp. 1, PP 1 | tentative | Lubricant, hydrocarbon resin emulsion, sunscreen (C) | Irritant, toxic to aquatic life (C) |
| 14130-06-4 | docosan-1-amine | HDPE 4, PS 5a | - | Hair cosmetics, heat transfer fluids (C) | - |
| 554-91-6 | gentiobiose | HDPE 4 | - | Biological molecule (C) | - |
| 20198-87-2 | N-benzyloctadecan-1-amine | HDPE 4 | yes (C) | Thermoplastic elastomer (C) | - |
| 510-75-8 | 12-hydroxy-11-methyl-6-methylidene-16-oxo-15-oxapentacyclo[9.3.2.15,8.01,10.02,8]heptadec-13-ene-9-carboxylic acid | HDPE 4 | - | Fungicide (C) | - |
| 119344-86-4 | 2-(dimethylamino)-2-[(4-methylphenyl)methyl]-1-(4-morpholin-4-ylphenyl)butan-1-one | PP 1 | yes (B) | UV light absorber, paints and coatings (C) | Reproductive toxicity, harmful to aquatic life (C) |

| CAS | Name | High abundance in sample | Plastic related | Use | Toxicity |
|---|---|---|---|---|---|
| 94108-97-1 | [2-[2,2-bis(prop-2-enoyloxymethyl)butoxymethyl]-2-(prop-2-enoyloxymethyl)butyl] prop-2-enoate | PP 1, PP 5b, PP 5a | yes (B) | Colorant, monomer (B) | Irritant (C). Not classified as of potential concern (B) |
| 1541-67-9 | lauryldiethanolamine | PP 3, PS 3 | yes (C) | Flame retardant, PUR-resin forming composition, propylene resin composition (C) | Irritant, toxic to aquatic life, genotoxic (C) |
| 1733-93-3 | 2-[2-dodecoxyethyl(2-hydroxyethyl)amino]ethanol | PP 3, PS 3 | tentative | Cosmetics, antistatic agent, surfactant (C) | - |
| 10213-78-2 | 2-[2-hydroxyethyl(octadecyl)amino]ethanol | PP 4 | yes (B) | Antistatic agent, emulsifying, lubricants, processing aids (B, C) | Irritant, corrosive, toxic to aquatic life (C). Persistant and bioaccumulative (B) |
| 52497-24-2 | 2-[2-hydroxyethyl(octadecyl)amino]ethyl octadecanoate | PP 4 | yes (B) | Antistatic agent, FCA (B, C) | Corrosive, irritant (C). Not classified as of potential concern (B) |
| 18924-66-8 | 2,2'-(Tetradecylimino)diethanol | PP 4 | yes (C) | Antistatic agent, PUR resinforming, composition, PS resin particles (B, C) | Corrosive, irritant (C). Not classified as of potential concern in (B). AR antagonist, ERα, PPARγ and PXR agonist (D) |
| 14174-09-5 | 2,5,8,11,18,21,24,27-octaoxatricyclo[26.4.0.012,17]dotriaconta-1(32),12,14,16,28,30-hexaene | PP 4 | tentative | Adhesive, polymers for use as anion exchange membranes (C) | Irritant (C) |
| 23681-60-9 | tris(4-anilinophenyl)methanol | PP 5a, PP 5b | - | Purification of triphenylmethane compounds (C) | - |

| CAS | Name | High abundance in sample | Plastic related | Use | Toxicity |
|---|---|---|---|---|---|
| 82799-44-8 | 2,4-diethylthioxanthen-9-one | PP 5a, PP 5b | yes (B) | Adhesive, colorant, filler, other processing aids (B) | Not classified as of potential concern (B) |
| 68966-33-6 | (4-aminophenyl)-bis(4-anilinophenyl)methanol | PP 5a, PP 5b | - | - | - |
| 4687-94-9 | [2-hydroxy-3-[4-[2-[4-(2-hydroxy-3-prop-2-enoyloxypropoxy)phenyl]propan-2-yl]phenoxy]propyl] prop-2-enoate | PP 5a, PP 5b | yes (B) | Monomer (B) | Irritant (C). Not classified as of potential concern (B) |
| 29649-48-7 | N-[5-[2-(2,5-dioxopyrrolidin-1-yl)ethyl-ethylamino]-2-[(4-nitrophenyl)diazenyl]phenyl]acetamide | PP 5a, PP 5b | - | - | - |
| - | 2-[[4-[(4-aminophenyl)-(4-anilinophenyl)methylidene]cyclohexa-2,5-dien-1-ylidene]amino]benzenesulfonic acid | PP 5b | - | - | - |
| 119313-12-1 | 2-benzyl-2-(dimethylamino)-1-(4-morpholin-4-ylphenyl)butan-1-one | PP 5a, PP 5b | yes (B) | Colorant, catalyst, filler, lubricant, other processing aids (B) | Persistent, bioaccumulate, CMR, reprotoxic, toxic to aquatic life (B). AR antagonist, ERα, PPARγ and PXR agonist (D) |
| 67989-24-6 | (3-hydroxy-2,2-dimethylpropyl) (Z)-octadec-9-enoate | PS 1 | tentative | Lubricants and lubricant additives (C) | - |
| 64050-16-4 | 2-[2,4-bis(2-methylbutan-2-yl)phenoxy]ethyl prop-2-enoate | PS 3 | - | - | - |

| CAS | Name | High abundance in sample | Plastic related | Use | Toxicity |
|---|---|---|---|---|---|
| 3386-18-3 | 2-[2-[2-[2-[2-[2-[2-[2-(2-hydroxyethoxy)ethoxy]ethoxy]ethoxy]ethoxy]ethoxy]ethoxy]ethoxy]ethanol | PS 4 | yes (C) | Processing aids and additives, coating (C) | Irritant (C) |
| 611-92-7 | 1,3-dimethyl-1,3-diphenylurea | PS 4 | - | - | Irritant, toxic to aquatic life (C) |
| 5617-32-3 | 2-[2-[2-[2-[2-[2-(2-hydroxyethoxy)ethoxy]ethoxy]ethoxy]ethoxy]ethoxy]ethanol | PS 4 | tentative | Processing aids and additives (C) | |
| 17598-96-8 | 2-[2-[2-[2-[2-[2-[2-[2-[2-[2-[2-(2-hydroxyethoxy)ethoxy]ethoxy]ethoxy]ethoxy]ethoxy]ethoxy]ethoxy]ethoxy]ethoxy]ethoxy]ethanol | PS 4 | - | Cosmetics (C) | - |
| 596-27-0 | 3,3-bis(4-hydroxy-3-methylphenyl)-2-benzofuran-1-one | PS 4 | - | - | - |
| 506-12-7 | heptadecanoic acid | PS 5a | yes (B) | Surfactant, lubricant, other processing aids (B, C) | Not classified as of potential concern (B) |
| 111-60-4 | 2-hydroxyethyl octadecanoate | PS 5a | yes (B) | Emulsifier, plasticizer, food additive (B, C) | Not classified as of potential concern (B) |
| 71989-20-3 | (2S)-5-amino-2-(9H-fluoren-9-ylmethoxycarbonylamino)-5-oxopentanoic acid | PS 5a | - | Peptide (C) | - |
| 69701-99-1 | (2-ethyl-2-methyl-1,3-dioxolan-4-yl)methyl prop-2-enoate | comp. 1 | tentative | Ink, adhesives (C) | Irritant (C) |
| 507-60-8 | [8,14-dihydroxy-10,13-dimethyl-17-(6-oxopyran-3-yl)-3-[3,4,5- | comp. 1 | - | rodenticide (C) | |

| CAS | Name | High abundance in sample | Plastic related | Use | Toxicity |
|---|---|---|---|---|---|
| | trihydroxy-6-(hydroxymethyl)oxan-2-yl]oxy-2,3,6,7,9,11,12,15,16,17-decahydro-1H cyclopenta[a]phenanthren-6-yl] acetate | | | | |
| 7441-32-9 | 3,6,13,16,23,26-hexaoxatetracyclo[26.2.2.28,11.218,21]hexatriaconta-1(31),8,10,18(34),19,21(33),28(32),29,35-nonaene-2,7,12,17,22,27-hexone | PET 1, PET 3, PET 4, PET 5, PET 6a | yes (B) | NIAS in PET (A) | - |
| 58-18-4 | (8R,9S,10R,13S,14S,17S)-17-hydroxy-10,13,17-trimethyl-2,6,7,8,9,11,12,14,15,16-decahydro-1H-cyclopenta[a]phenanthren-3-one | PET 2 | - | Drug, steroid (C) | May cause cancer and reproductive toxicity (C). AR antagonist, ERα, PPARγ and PXR agonist (D) |
| - | (8S,9S,10S,13S,14S)-10,13-dimethyl-4,5,6,7,8,9,11,12,14,15,16,17-dodecahydro-3H-cyclopenta[a]phenanthren-1-ol | PET 2 | - | Pharmaceutical (C) | - |
| 62488-56-6 | nona-2,4-dien-1-ol | PET 2 | - | Flavoring agent in food, ink (C) | - |
| 69686-05-1 | 1,8-dihydroxy-3-methyl-6-[(3R,4R,5R,6S)-3,4,5-trihydroxy-6- | PET 4 | - | - | - |

| CAS | Name | High abundance in sample | Plastic related | Use | Toxicity |
|---|---|---|---|---|---|
| | methyloxan-2-yl]oxyanthracene-9,10-dione | | | | |
| 10287-53-3 | ethyl 4-(dimethylamino)benzoate | PET 5 | yes (B) | Colorant, filler, other processing aids (B) | Persistent and bioaccumulative, carcinogenic, mutagenic and reprotoxic, toxic to aquatic life (B, C) ERα, PPARγ and PXR agonist (D) |
| 84-11-7 | phenanthrene-9,10-dione | PET 5 | tentative | Used in prod. of dyes, herbicides, pharmaceuticals, UV curable coatings and adhesives (C) | Irritant, cytotoxic (C) |
| 18595-18-1 | methyl 3-amino-4-methylbenzoate | PET 5 | - | - | Irritant (C) |
| 18814-21-6 | 1-[3,4-dihydroxy-5-(hydroxymethyl)oxolan-2-yl]-5-fluoropyrimidine-2,4-dione | PET 5 | - | Pharmaceutical (C) | - |
| 3147-75-9 | Octrizole | PUR 1 | yes (B) | UV absorber, antioxidant, UV light stabilizer in plastic, used in cosmetics, used in food contact substances (C) | Irritant, environmental hazard (C), not classified as of potential concern in (B), AR antagonist, ERα and PXR agonist (D) |
| 115-86-6 | Triphenyl phosphate | PUR 1, PUR 2 | yes (B) | Plasticizer, other processing aids, FCA (A, C) | Environmental hazard, toxic to aquatic life, long-term hazard (C), Bioaccumulative, persistent, aquatic toxicity and EDC (B), AR antagonist, ERα, PPARγ and PXR agonist (D) |
| 41556-26-7 | bis(1,2,2,6,6-pentamethylpiperidin-4-yl) decanedioate | PUR 2 | yes (B) | UV light absorber, stabilizer (C) | Toxic to aquatic life, irritant, persistant and bioaccumulative (B, C) |

| CAS | Name | High abundance in sample | Plastic related | Use | Toxicity |
|---|---|---|---|---|---|
| 82919-37-7 | 1-O-methyl 10-O-(1,2,2,6,6-pentamethylpiperidin-4-yl) decanedioate | PUR 2 | yes (B) | Antioxidant, stabilizer (C) | Toxic to aquatic life, irritant, persistant and bioaccumulative (B, C) |
| 57834-33-0 | ethyl 4-[(N-methylanilino)methylideneamino]benzoate | PUR 2 | yes (B) | Colorant, filler, light stabilizer, other processing aids (B) | Toxic to aquatic life (long term hazard), harmful if swallowed, irritant (C), Not classified as of potential concern (B) |
| 67845-92-5 | 8-(2,2,6,6-tetramethylpiperidine-4-carbonyl)oxyoctyl 2,2,6,6-tetramethylpiperidine-4-carboxylate | PUR 2 | yes (B) | Polyester compositions (C) | - |
| 5834-63-9 | 1,8,15,22-tetrazacyclooctacosane-2,9,16,23-tetrone | comp. 2 | yes (A, B) | | - |
| 4266-66-4 | 1,8-diazacyclotetradecane-2,7-dione | comp. 2 | yes (B) | Flame retardant, stabilizer (C) | Not classified as of potential concern (B) |
| 96507-76-5 | 2-O-decyl 1-O-nonyl benzene-1,2-dicarboxylate | PVC 2 | yes (C) | Plasticizer (C) | - |
| - | [(3R,4S,5S,6R)-3,4,5-trihydroxy-6-(hydroxymethyl)oxan-2-yl] (Z)-octadec-9-enoate | PVC 3 | - | Alkyl glycoside-based micellar thickeners for surfactant systems (C) | - |
| 56149-06-5 | 2,3-diacetyloxypropyl docosanoate | PVC 3 | - | - | - |
| 92739-54-3 | 10-dodecoxy-10-oxodecanoic acid | PVC 4 | tentative | In inks containing crystalline polyesters (C) | - |

#### S2.4 Limitations of the chemical analysis and identifications.

It should be noted that the number of chemicals reported here is not comprehensive as the measurements were only performed using positive ionization mode. This means that negatively ionizable compounds were not detected, possibly leading to an underestimation of the actual number of chemicals present. Yet, the largest fraction of plastic related chemicals is positively ionizable.[13] One-third of the samples analyzed had previous food content, which can alter the chemical composition to varying extents, as demonstrated by a comparison of three products analyzed both with and without previous food content. This could result in an overestimation; however, no general trend of higher numbers was observed in samples with previous food content. Furthermore, the four samples with the highest number of features did not have previous food content.

The low identification rate, as achieved in this study, is a known challenge, especially with liquid chromatographic methods. The scarcity of publicly available information on plastic chemicals paired with technical challenges of identification of LC-MS data hinders a more comprehensive identification.[14] While there is much uncertainty regarding the use of chemicals in plastics, about 55% of the top ten most abundant features in our samples were not clearly related to plastics based on the sources we consulted. These chemicals may have originated from the food content, be formed or introduced during plastic production (e.g., NIAS), are used in plastic production but not known to the public, or simply be misidentified.[5,15,16]

In another study which used a nontarget liquid chromatography coupled to high-resolution mass spectrometry,[17] 76% of the identified chemicals in food packaging were classified as NIAS. In accordance with this, Geueke et al. reported that 65% of chemicals in food contact materials, including plastics, are not listed for their use in food contact applications.[18] Nonetheless, 45% of the ten most abundant features were identified as chemicals commonly used in plastics, indicating the suitability of this nontarget approach.

#### S2.5 Receptor activity of the tentatively identified chemicals retrieved from ToxCast

For prioritization of the chemicals, we used a ranking system to identify the most abundant chemicals in the samples as a proxy for their concentration and their potency of receptor activity according to ToxCast (Table S9). To do so, we calculated the ratio of the chemical's $AC_{50}$ over its abundance. Within the highest-ranking chemicals, we found the plastic related chemicals triphenyl phosphate (CAS 115-86-6), which is the highest ranking PXR ($AC_{50}$ = 0.7 µM) and PPARγ ($AC_{50}$ = 0.003 µM) agonist and was detected in ten samples made of six polymers or octrizole (CAS 3147-75-9, 2nd rank for PXR, $AC_{50}$ = 1.0 µM) and tributyl 2-acetyloxypropane1,2,3-tricarboxylate (CAS 77-90-7, 3rd rank for PPARγ, $AC_{50}$ = 7.4 µM). The ERα agonist with the lowest $AC_{50}$/abundance ratio is 4-dodecyl phenol (CAS 104-43-8, $AC_{50}$ = 0.08 µM) which was detected in six FCAs (LDPE 8, PUR 1, PVC 3, 4, and comp. 1 and 2) and is an additive in lubricants and fuels. The highest-ranking anti-androgenic compound was cyclohexanone oxime (CAS 100-64-1, $AC_{50}$ = 0.7 µM), a plastic-associated chemical detected in three FCAs (PUR 1, PVC 4 and comp. 2). Further, looking at high-potency plastic chemicals which were detected in more than six samples of different polymers, the plasticizer triethylene glycol (CAS 112-27-6, PXR $AC_{50}$ = 0.11 µM) and tetraethylene glycol, (CAS 112-27-6, PPARγ $AC_{50}$ = 0.06 µM) are notable. These compounds were detected in ten FCAs made of four polymers (LDPE 2, 4, 7, HDPE 3, comp. 1, PET 1, 5, PP 4, PS 4 and 5b) and in eight FCAs covering four polymers and the composites (LDPE 4, 7, 8, PP 5b, PS 5b, PVC 4, comp. 1 and 2), respectively. Both tetraethylene glycol and triethylene glycol were also identified as AR antagonists.

Table S9. PXR, PPARγ, ERα agonists and AR antagonists tentatively identified in the plastic extracts. Compounds ranked based on $AC_{50}$/peak area per receptor. First 20 compounds per receptor are listed for others see Table S15.

| CAS | Rank | $AC_{50}$ [µM] | Highest peak | Ratio $AC_{50}$/peak | Detected in samples |
|---|---|---|---|---|---|
| **PXR** | | | | | |
| 115-86-6 | 1 | 0.70 | 542944 | 1.30E-06 | PUR_1, PUR_2, PVC_2, LDPE_4, LDPE_7, comp_1, PET_4, PET_5, PP_5b, PS_4 |
| 3147-75-9 | 2 | 1.04 | 607970 | 1.71E-06 | PUR_1, PUR_2, PET_2 |
| 77-90-7 | 3 | 0.85 | 184096 | 4.63E-06 | comp_2, LDPE_2, PS_5b |
| 791-28-6 | 4 | 0.70 | 132896 | 5.28E-06 | PUR_1, PUR_2, comp_2, PP_5b, PP_5a |
| 59227-89-3 | 5 | 1.41 | 191304 | 7.36E-06 | PUR_1, comp_2, PVC_3, PVC_4, LDPE_8, LDPE_5, LDPE_7, LDPE_2, LDPE_3, comp_1 |
| 112-27-6 | 6 | 0.11 | 2626 | 4.37E-05 | LDPE_4, LDPE_7, HDPE_3, LDPE_2, comp_1, PET_1, PET_5, PP_4, PS_4, PS_5b |
| 104-43-8 | 7 | 1.45 | 31244 | 4.64E-05 | PUR_1, comp_2, PVC_3, PVC_4, LDPE_8, comp_1 |
| 119313-12-1 | 8 | 1.63 | 31445 | 5.18E-05 | LDPE_4, LDPE_7, PP_1, PP_5b, PP_5a |
| 18924-66-8 | 9 | 2.22 | 39595 | 5.60E-05 | LDPE_7, PP_4 |
| 4986-89-4 | 10 | 0.64 | 6559 | 9.81E-05 | PP_1, PP_5b, PP_5a, PS_1 |
| 124-07-2 | 11 | 0.09 | 548 | 1.68E-04 | PUR_1, PUR_2, PVC_1, PVC_3, PVC_4 |
| 3179-80-4 | 12 | 4.15 | 15416 | 2.69E-04 | comp_2 |
| 68-26-8 | 13 | 0.30 | 952 | 3.14E-04 | LDPE_4, LDPE_3, comp_1, |
| 94-62-2 | 14 | 1.26 | 1989 | 6.34E-04 | LDPE_4, LDPE_7, comp_1, PET_6b, |
| 75330-75-5 | 15 | 1.66 | 2163 | 7.67E-04 | PVC_1 |
| 117-81-7 | 16 | 9.14 | 10032 | 9.11E-04 | PVC_2 |
| 78-51-3 | 17 | 0.99 | 984 | 1.00E-03 | LDPE_4, |
| 55179-31-2 | 18 | 1.16 | 915 | 1.27E-03 | PUR_2, PVC_4, |
| 42978-66-5 | 19 | 2.83 | 1926 | 1.47E-03 | LDPE_8, comp_1, PP_5b, PP_5a, |
| 6707-60-4 | 20 | 33.24 | 17349 | 1.92E-03 | PUR_1, PUR_2, comp_2, PVC_3, PVC_4, LDPE_8, comp_1, PET_2, PS_5b, |
| **PPARγ** | | | | | |
| 115-86-6 | 1 | 0.003 | 542944 | 5.55E-09 | PUR_1, PUR_2, PVC_2, LDPE_4, LDPE_7, comp_1, PET_4, PET_5, PP_5b, PS_4 |
| 87-62-7 | 2 | 0.014 | 5892 | 2.43E-06 | PUR_2, PVC_3, PVC_4 |
| 77-90-7 | 3 | 7.39 | 184096 | 4.01E-05 | comp_2, LDPE_2, PS_5b, |
| 59227-89-3 | 4 | 7.87 | 191304 | 4.12E-05 | PUR_1, comp_2, PVC_3, PVC_4, LDPE_8, LDPE_5, LDPE_7, LDPE_2, LDPE_3, comp_1, |
| 112-60-7 | 5 | 0.06 | 745 | 8.40E-05 | comp_2, PVC_4, LDPE_8, LDPE_4, LDPE_7, comp_1, PP_5b, PS_5b, |

| CAS | Rank | $AC_{50}$ [µM] | Highest peak | Ratio $AC_{50}$/peak | Detected in samples |
|---|---|---|---|---|---|
| 4986-89-4 | 6 | 1.78 | 6559 | 2.71E-04 | PP_1, PP_5b, PP_5a, PS_1, |
| | | | | | PUR_1, PUR_2, comp_2, PVC_3, PVC_4, LDPE_4, |
| 142-18-7 | 7 | 13.01 | 25094 | 5.19E-04 | PET_2, |
| 105-95-3 | 8 | 92.22 | 99278 | 9.29E-04 | PVC_3, |
| 119313-12-1 | 9 | 42.25 | 31445 | 1.34E-03 | LDPE_4, LDPE_7, PP_1, PP_5b, PP_5a, |
| 94-09-7 | 10 | 40.96 | 17969 | 2.28E-03 | PUR_1, PUR_2, PVC_3, PVC_4, LDPE_4, |
| 117-81-7 | 11 | 25.99 | 10032 | 2.59E-03 | PVC_2 |
| | | | | | comp_2, PVC_3, PVC_4, LDPE_4, LDPE_5, |
| 143-07-7 | 12 | 10.05 | 3872 | 2.60E-03 | LDPE_7, LDPE_2, LDPE_3, comp_1, |
| 58846-77-8 | 13 | 21.51 | 8029 | 2.68E-03 | comp_2, PVC_1, PVC_3, PVC_4, LDPE_7, PS_4, |
| 104-43-8 | 14 | 89.04 | 31244 | 2.85E-03 | PUR_1, comp_2, PVC_3, PVC_4, LDPE_8, comp_1, |
| | | | | | PUR_1, PUR_2, comp_2, PVC_3, PVC_4, LDPE_8, |
| 6707-60-4 | 15 | 63.93 | 17349 | 3.68E-03 | comp_1, PET_2, PS_5b, |
| 1541-67-9 | 16 | 66.65 | 13092 | 5.09E-03 | LDPE_7, PP_3, PS_3, |
| 3179-80-4 | 17 | 85.37 | 15416 | 5.54E-03 | comp_2, |
| 8004-87-3 | 18 | 9.38 | 1128 | 8.32E-03 | PP_5b, PP_5a, |
| 42978-66-5 | 19 | 17.29 | 1926 | 8.98E-03 | LDPE_8, comp_1, PP_5b, PP_5a, |
| 16219-75-3 | 20 | 24.07 | 2636 | 9.13E-03 | PVC_1, PVC_3, PVC_4, |
| **ERα** | | | | | |
| 104-43-8 | 1 | 0.08 | 31244 | 2.70E-06 | PUR_1, comp_2, PVC_3, PVC_4, LDPE_8, comp_1, |
| 521-17-5 | 2 | 0.08 | 11309 | 7.12E-06 | PUR_1, PVC_1, PVC_2, PVC_3, PVC_4, |
| | | | | | PUR_1, PUR_2, PVC_2, LDPE_4, LDPE_7, comp_1, |
| 115-86-6 | 3 | 7.98 | 542944 | 1.47E-05 | PET_4, PET_5, PP_5b, PS_4 |
| 123-34-2 | 4 | 0.06 | 3839 | 1.67E-05 | comp_2, PVC_3, PVC_4, |
| 119313-12-1 | 5 | 0.96 | 31445 | 3.06E-05 | LDPE_4, LDPE_7, PP_1, PP_5b, PP_5a, |
| 3147-75-9 | 6 | 31.55 | 607970 | 5.19E-05 | PUR_1, PUR_2, PET_2, |
| 25564-22-1 | 7 | 0.06 | 205 | 3.01E-04 | LDPE_7, |
| 57-11-4 | 8 | 2.30 | 2043 | 1.12E-03 | PVC_3, PVC_4, LDPE_8, |
| 75330-75-5 | 9 | 7.09 | 2163 | 3.28E-03 | PVC_1, |
| 102-76-1 | 10 | 39.31 | 7900 | 4.98E-03 | PUR_1, PVC_3, PVC_4, |
| 67-73-2 | 11 | 4.83 | 864 | 5.59E-03 | PP_1, PP_5b, PP_5a, |
| 117-81-7 | 12 | 59.54 | 10032 | 5.94E-03 | PVC_2 |
| | | | | | PUR_1, PVC_2, comp_1, PP_5a, PS_4, PS_5a, |
| 65-85-0 | 13 | 69.32 | 11145 | 6.22E-03 | PS_5b, |
| 135-19-3 | 14 | 5.70 | 767 | 7.43E-03 | PP_1, PS_3, PS_4, PS_5a, PS_5b, |
| 87-51-4 | 15 | 13.96 | 1843 | 7.58E-03 | comp_2, |

| CAS | Rank | $AC_{50}$ [µM] | Highest peak | Ratio $AC_{50}$/peak | Detected in samples |
|---|---|---|---|---|---|
| 4940-11-8 | 16 | 8.83 | 976 | 9.05E-03 | PUR_1, PUR_2, comp_2, PVC_4, |
| 620-40-6 | 17 | 3.98 | 428 | 9.29E-03 | PUR_1, |
| 1224-92-6 | 18 | 18.53 | 1687 | 1.10E-02 | LDPE_4, LDPE_5, LDPE_6, LDPE_7, LDPE_2, LDPE_3, comp_1, PP_1, PP_5b, PP_5a, PS_4, |
| 21145-77-7 | 19 | 4.80 | 299 | 1.60E-02 | PVC_3, PVC_4, |
| 57-11-4 | 20 | 2.30 | 143 | 1.61E-02 | PE_LD_44, LDPE_4, LDPE_6, PE_LP_10, PE_LP_15, PS_4, PS41a |
| **AR** | | | | | |
| 100-64-1 | 1 | 0.69 | 19747 | 3.50E-05 | PUR_1, comp_2, PVC_4, |
| 115-86-6 | 2 | 25.28 | 542944 | 4.66E-05 | PUR_1, PUR_2, PVC_2, LDPE_4, LDPE_7, comp_1, PET_4, PET_5, PP_5b, PS_4 |
| 3147-75-9 | 3 | 31.39 | 607970 | 5.16E-05 | PUR_1, PUR_2, PET_2, |
| 59227-89-3 | 4 | 42.55 | 191304 | 2.22E-04 | PUR_1, comp_2, PVC_3, PVC_4, LDPE_8, LDPE_5, LDPE_7, LDPE_2, LDPE_3, comp_1, |
| 87-62-7 | 5 | 1.62 | 5892 | 2.75E-04 | PUR_2, PVC_3, PVC_4 |
| 18924-66-8 | 6 | 12.48 | 39595 | 3.15E-04 | LDPE_7, PP_4, |
| 94-09-7 | 7 | 6.07 | 17969 | 3.38E-04 | PUR_1, PUR_2, PVC_3, PVC_4, LDPE_4, |
| 4986-89-4 | 8 | 3.15 | 6559 | 4.80E-04 | PP_1, PP_5b, PP_5a, PS_1, |
| 104-43-8 | 9 | 16.22 | 31244 | 5.19E-04 | PUR_1, comp_2, PVC_3, PVC_4, LDPE_8, comp_1, |
| 99-96-7 | 10 | 0.60 | 1102 | 5.48E-04 | PVC_1, PVC_2, |
| 75-07-0 | 11 | 1.15 | 1506 | 7.66E-04 | PVC_3, PVC_4, |
| 96-41-3 | 12 | 0.27 | 305 | 8.72E-04 | comp_2, PVC_3, |
| 78-51-3 | 13 | 0.89 | 984 | 9.00E-04 | LDPE_4, |
| 495-18-1 | 14 | 40.58 | 40617 | 9.99E-04 | PUR_1, PUR_2, |
| 119313-12-1 | 15 | 31.86 | 31445 | 1.01E-03 | LDPE_4, LDPE_7, PP_1, PP_5b, PP_5a, |
| 3055-99-0 | 16 | 5.86 | 5544 | 1.06E-03 | comp_2, |
| 3179-80-4 | 17 | 16.36 | 15416 | 1.06E-03 | comp_2, |
| 8004-87-3 | 18 | 1.38 | 1128 | 1.23E-03 | PP_5b, PP_5a, |
| 75330-75-5 | 19 | 3.57 | 2163 | 1.65E-03 | PVC_1, |
| 1541-67-9 | 20 | 28.60 | 13092 | 2.18E-03 | LDPE_7, PP_3, PS_3, |

Table S10. Number of known, active chemicals detected in the samples compared to the experimentally derived effect concentrations.

| Sample | PXR agonists/ antagonists | $EC_{20}$ [mg well$^{-1}$] | PPARg agonists/ antagonists | $EC_{20}$ [mg well$^{-1}$] | ERa agonists/ antagonists | $EC_{20}$ [mg well$^{-1}$] | AR antagonists/ agonists | $EC_{50}$ [mg well$^{-1}$] |
|---|---|---|---|---|---|---|---|---|
| HDPE 1 | 0/0 | 0.26 | 0/0 | 0 | 0/0 | 0 | 0/0 | 0 |
| HDPE 2 | 0/0 | 0 | 0/0 | 0 | 0/0 | 0 | 0/0 | 0 |
| HDPE 3 | 3/0 | 0.15 | 1/0 | 1.03 | 2/0 | 0 | 1/0 | 0 |
| HDPE 4 | 1/0 | 0 | 0/0 | 0 | 0/1 | 0 | 2/0 | 0 |
| LDPE 1 | 1/0 | 0 | 1/0 | 0 | 0/0 | 0 | 0/0 | 0 |
| LDPE 2 | 9/0 | 0.06 | 2/0 | 0.46 | 4/1 | 0 | 3/1 | 0.75 |
| LDPE 3 | 8/0 | 0.12 | 4/0 | 0.47 | 3/1 | 0 | 3/1 | 0.52 |
| LDPE 4 | 20/0 | 0.13 | 6/1 | 1.10 | 10/4 | 0 | 11/4 | 0 |
| LDPE 5 | 5/0 | 0.06 | 2/0 | 0.21 | 1/1 | 0 | 1/0 | 0.41 |
| LDPE 6 | 4/0 | 0.11 | 0/0 | 0.42 | 2/2 | 0 | 4/1 | 1.06 |
| LDPE 7 | 25/0 | 0.07 | 7/1 | 0.59 | 15/4 | 0.15 | 9/4 | 0.73 |
| LDPE 8 | 8/0 | 0 | 5/1 | 0.37 | 6/1 | 0 | 5/1 | 0 |
| PET 1 | 4/0 | 0.21 | 1/0 | 0 | 0/1 | 0 | 5/1 | 0 |
| PET 2 | 8/1 | 0.19 | 4/0 | 0 | 6/0 | 0 | 2/1 | 0 |
| PET 3 | 0/0 | 0.73 | 0/0 | 0 | 0/0 | 0 | 0/0 | 0 |
| PET 4 | 2/0 | 0 | 3/1 | 0 | 1/0 | 0 | 2/0 | 0 |
| PET 5 | 4/0 | 0.78 | 2/1 | 0 | 3/0 | 0 | 4/0 | 0 |
| PET 6a | 0/0 | 0.54 | 0/0 | 0 | 0/0 | 0 | 0/0 | 0 |
| PET 6b | 2/0 | 0.15 | 0/0 | 0 | 1/0 | 0 | 1/0 | 0 |
| PP 1 | 13/0 | 0.25 | 6/0 | 0 | 8/4 | 0 | 5/0 | 0 |
| PP 2 | 0/0 | 1 | 0/0 | 0 | 1/0 | 0 | 0/0 | 0 |
| PP 3 | 2/0 | 1 | 2/0 | 0 | 0/1 | 0 | 2/0 | 0 |
| PP 4 | 4/0 | 0.37 | 4/0 | 0 | 3/2 | 0 | 4/1 | 0 |
| PP 5a | 10/0 | 0.16 | 6/1 | 0 | 6/4 | 1 | 7/4 | 0 |
| PP 5b | 6/0 | 0.22 | 5/0 | 0 | 4/3 | 0.00 | 5/4 | 0 |
| PS 1 | 2/0 | 0.39 | 1/0 | 0 | 0/1 | 0.81 | 2/0 | 0 |
| PS 2 | 0/0 | 0.60 | 0/0 | 0 | 0/0 | 0 | 0/0 | 0 |
| PS 3 | 4/0 | 0.48 | 3/0 | 0 | 2/2 | 0.90 | 3/0 | 0 |
| PS 4 | 13/1 | 0.15 | 5/1 | 0 | 9/4 | 0.65 | 10/3 | 0 |
| PS 5a | 5/0 | 0.16 | 3/0 | 0 | 3/1 | 0.21 | 3/0 | 1.13 |
| PS 5b | 7/1 | 0.07 | 4/0 | 0 | 5/1 | 0.09 | 4/0 | 0.67 |
| PUR 1 | 28/2 | 0 | 6/2 | 0 | 18/3 | 0 | 15/9 | 0 |
| PUR 2 | 30/1 | 0.01 | 7/3 | 0 | 14/5 | 0 | 18/7 | 0 |
| PVC 1 | 10/0 | 0.11 | 2/0 | 0.17 | 6/0 | 0 | 4/3 | 0.26 |
| PVC 2 | 6/0 | 0.51 | 5/0 | 1.14 | 6/0 | 0 | 5/2 | 0 |
| PVC 3 | 26/4 | 0.15 | 11/4 | 0.58 | 15/6 | 0 | 20/7 | 0 |
| PVC 4 | 26/6 | 0.01 | 10/3 | 0.15 | 16/10 | 0 | 22/7 | 0.29 |
| comp. 1 | 22/1 | 0.11 | 10/1 | 0.66 | 14/2 | 0 | 8/5 | 0 |
| comp. 2 | 31/6 | 0.01 | 10/2 | 0.53 | 12/8 | 0 | 16/5 | 0.88 |

#### S2.6 Predictors of receptor activity

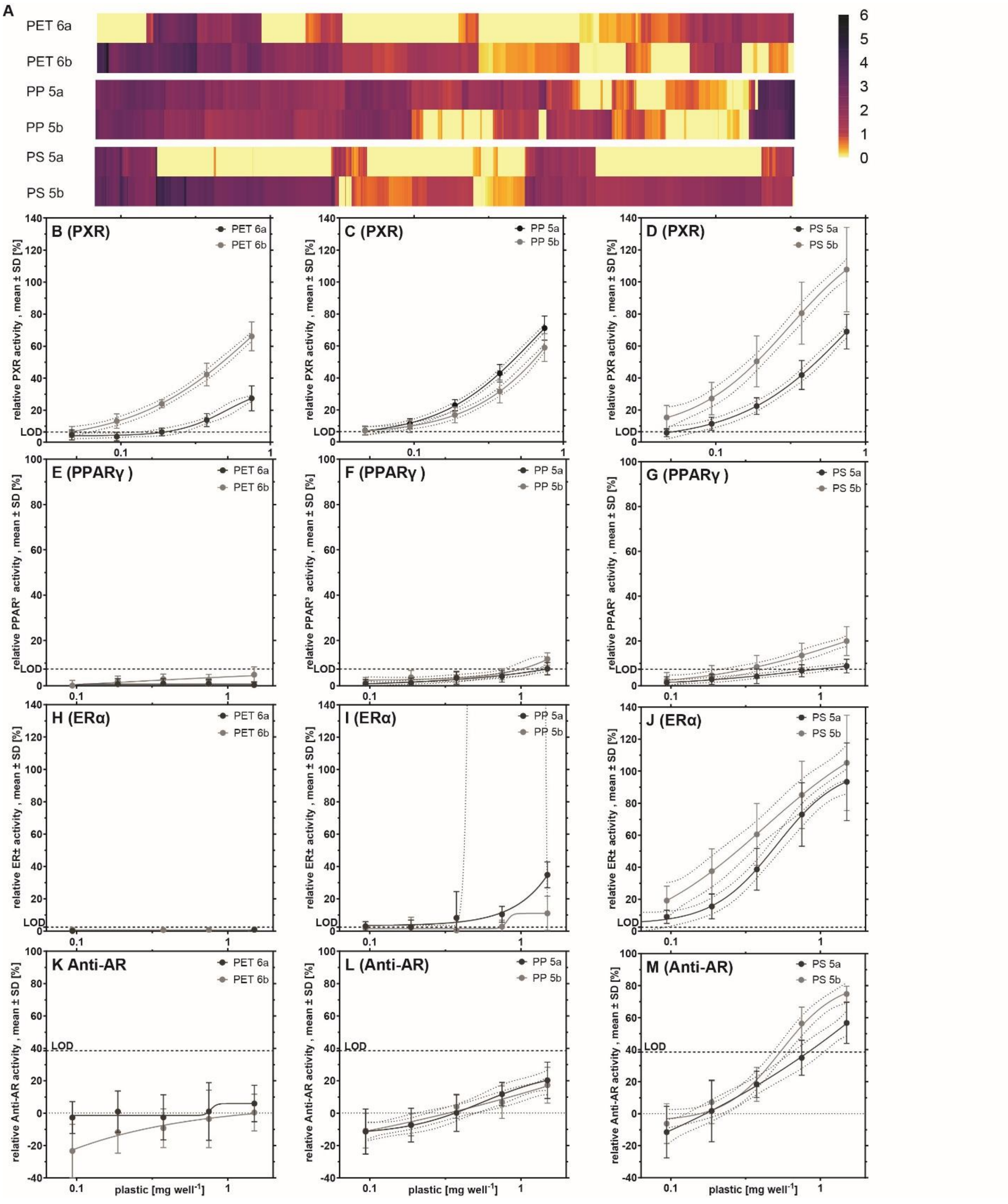


Figure S5. Comparison of samples without (a) and with (b) previous food content on chemical composition (A) and (B–M) receptor activity. Bioassay data are derived from at least three independent experiments, with four technical replicates per concentration (n ≥ 12).

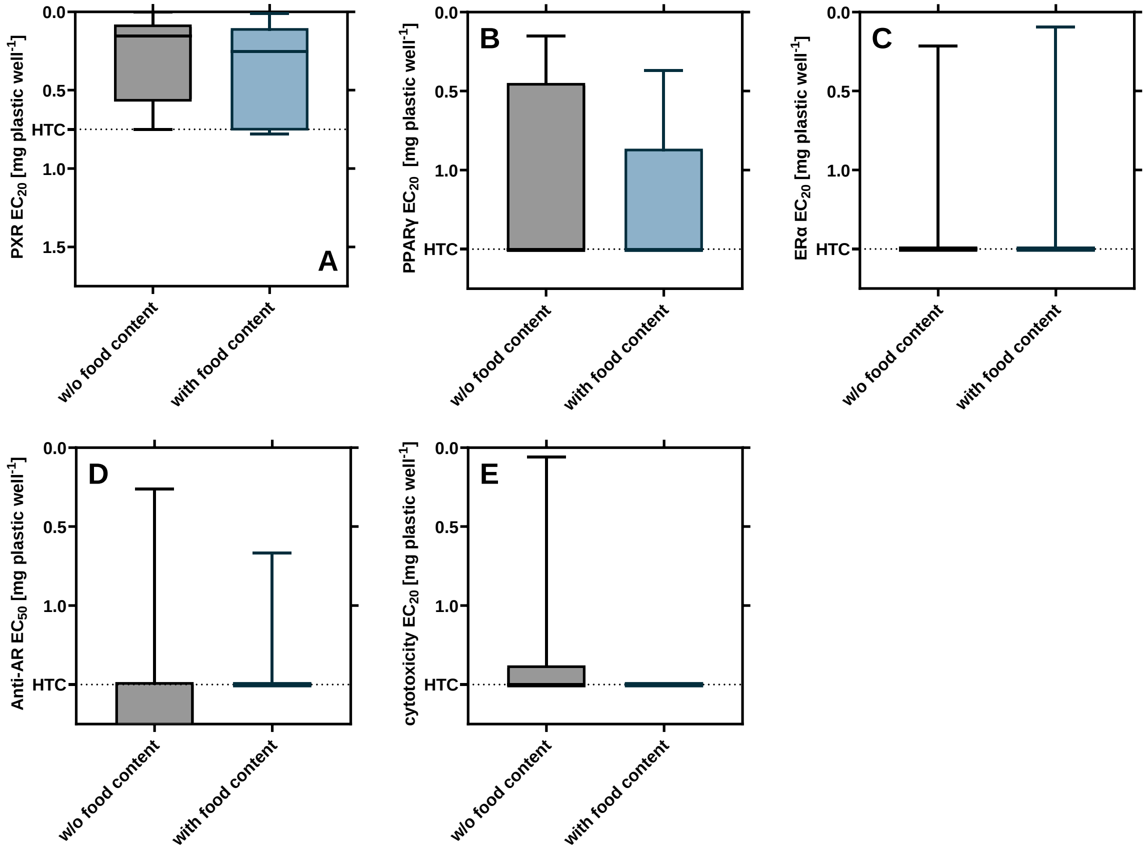


Figure S6**.** Effect of previous food content on receptor activity and cytotoxicity. $EC_{20/50}$ values calculated from a minimum of three independent experiments with four technical replicates per concentration ($n \geq 12$). Mann-Whitney tests do not indicate significant differences ($p > 0.05$).

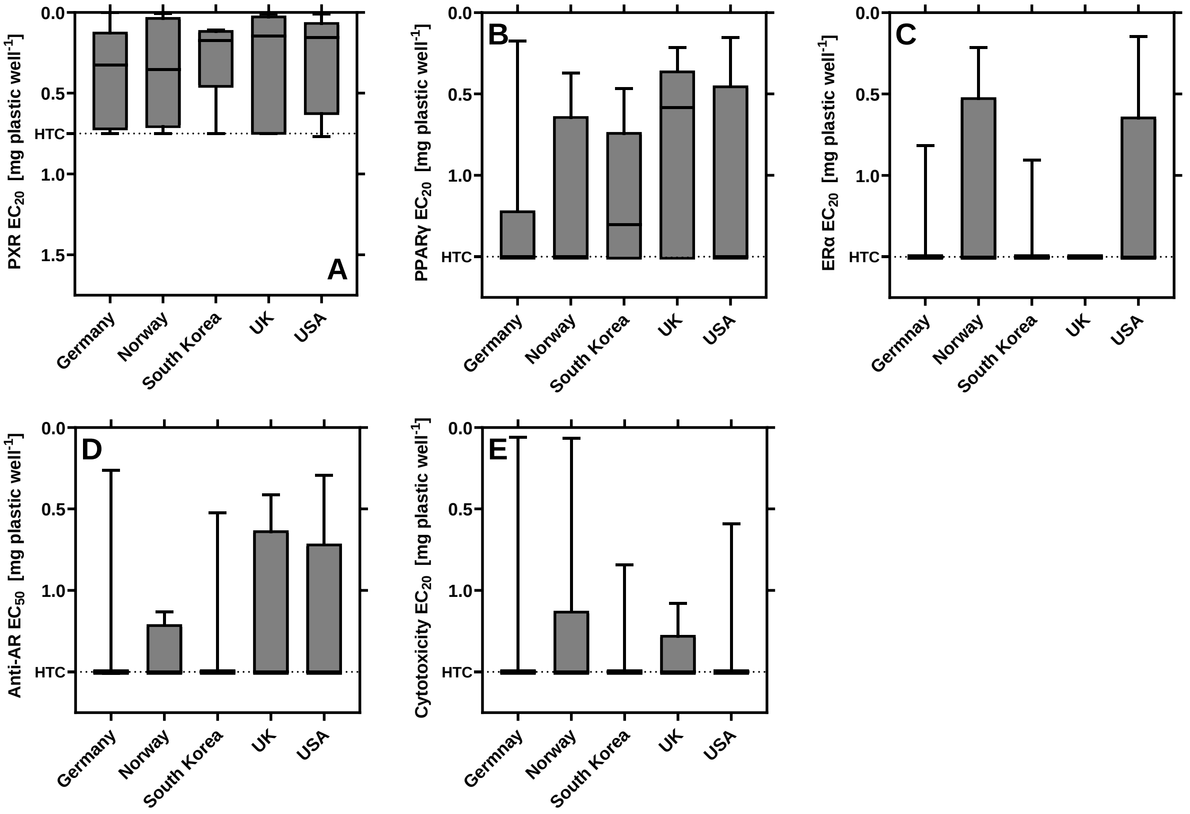


Figure S7. Impact of country of origin on receptor activity and cytotoxicity. $EC_{20/50}$ values calculated from a minimum of three independent experiments with four technical replicates per concentration ($n \geq 12$). Kruskal Wallis tests and Dunn's multiple comparison tests do not indicate significant differences ($p > 0.05$).

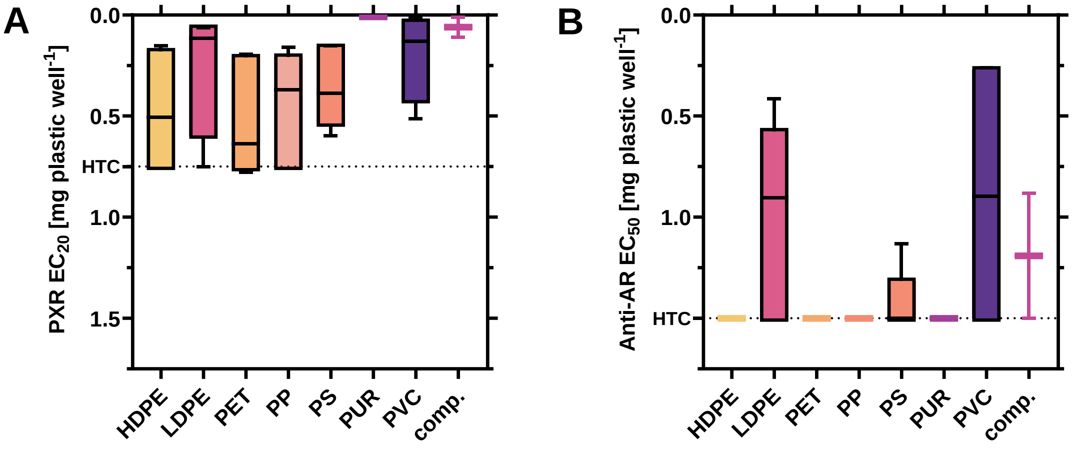


Figure S8. Impact of the polymer type on receptor activity at (A) PXR, (B) Anti-AR. $EC_{20/50}$ calculated from at least three independent experiments with four technical replicates per concentration ($n \geq 12$). Kruskal-Wallis tests with Dunn's multiple comparison tests do not indicate significant differences ($p > 0.05$).

#### S2.7 Partial least squares regression

Table S11**.** Performance of the PLS regression of chemical features (occurrence and abundance) and receptor activity (normalized EC20/50 values) after iterative filtering steps excluding features with VIP <0.8. PLS regression with food contact articles made of PE, PET, PP and PS. Calculated (Cal), cross-validated (CV), cumulative explained variance (cumexpvar) of the chemical features (X) and receptor activity (Y), determination coefficient ($R^2$), root mean squared errors (RMSE), slope for predicted vs. measured values (slope), bias for prediction vs. measured values (bias), ratio of standard deviation of response values to standard error of prediction (RPD).

| Model | N selected components | N features | | X cumexpvar | Y cumexpvar | $R^2$ | RMSE | Slope | Bias | RPD |
|---|---|---|---|---|---|---|---|---|---|---|
| **PXR** | | | | | | | | | | |
| 0 | 3 | 2993 | Cal | 46.0 | 76.9 | 0.77 | 17.3 | 0.769 | 0 | 2.11 |
| | | | Cv | NA | NA | 0.38 | 28.3 | 0.565 | -1.44 | 1.29 |
| 1 | 4 | 2147 | Cal | 69.89 | 88.93 | 0.89 | 12.0 | 0.889 | 0 | 3.05 |
| | | | Cv | NA | NA | 0.62 | 22.2 | 0.766 | -0.34 | 1.65 |
| 2 | 4 | 1761 | Cal | 67.80 | 89.99 | 0.90 | 11.4 | 0.9 | 0 | 3.21 |
| | | | Cv | NA | NA | 0.62 | 22.3 | 0.772 | 0.40 | 1.64 |
| 3 | 4 | 1583 | Cal | 65.56 | 90.39 | 0.90 | 11.2 | 0.904 | 0 | 3.28 |
| | | | Cv | NA | NA | 0.64 | 21.7 | 0.791 | 0.21 | 1.68 |
| 4 | 4 | 1544 | Cal | 63.03 | 90.54 | 0.91 | 11.1 | 0.905 | 0 | 3.3 |
| | | | Cv | NA | NA | 0.64 | 21.5 | 0.798 | 0.25 | 1.7 |
| 5 | 4 | 1536 | Cal | 62.39 | 90.57 | 0.91 | 11.1 | 0.906 | 0 | 3.31 |
| | | | Cv | NA | NA | 0.64 | 21.5 | 0.798 | 0.28 | 1.7 |
| **6** | **4** | **1533** | **Cal** | **62.25** | **90.58** | **0.91** | **11.1** | **0.906** | **0** | **3.31** |
| | | | **Cv** | **NA** | **NA** | **0.64** | **21.5** | **0.798** | **0.28** | **1.7** |
| **PPARγ** | | | | | | | | | | |
| 0 | 1 | 2993 | Cal | 25.53 | 69.80 | 0.70 | 17.4 | 0.698 | 0 | 1.85 |
| | | | Cv | NA | NA | 0.58 | 20.6 | 0.612 | -0.91 | 1.56 |
| 1 | 1 | 1232 | Cal | 81.48 | 69.13 | 0.69 | 17.6 | 0.691 | 0 | 1.83 |
| | | | Cv | NA | NA | 0.62 | 19.5 | 0.696 | -0.73 | 1.65 |
| 2 | 2 | 836 | Cal | 86.88 | 85.11 | 0.85 | 12.2 | 0.851 | 0 | 2.63 |
| | | | Cv | NA | NA | 0.68 | 17.8 | 0.765 | 0.70 | 1.81 |
| 3 | 2 | 756 | Cal | 85.00 | 85.45 | 0.86 | 12.1 | 0.855 | 0 | 2.66 |
| | | | Cv | NA | NA | 0.71 | 17.0 | 0.772 | 0.73 | 1.89 |
| 4 | 2 | 734 | Cal | 84.51 | 85.47 | 0.86 | 12.1 | 0.855 | 0 | 2.67 |
| | | | Cv | NA | NA | 0.72 | 16.8 | 0.772 | 0.75 | 1.91 |
| 5 | 2 | 731 | Cal | 84.20 | 85.48 | 0.86 | 12.0 | 0.855 | 0 | 2.67 |
| | | | Cv | NA | NA | 0.72 | 16.8 | 0.773 | 0.75 | 1.92 |
| **6** | **2** | **729** | **Cal** | **84.23** | **85.48** | **0.86** | **12.0** | **0.855** | **0** | **2.67** |
| | | | **Cv** | **NA** | **NA** | **0.72** | **16.8** | **0.773** | **0.75** | **1.92** |

| Model | N selected components | N features | | X cumexpvar | Y cumexpvar | $R^2$ | RMSE | Slope | Bias | RPD |
|---|---|---|---|---|---|---|---|---|---|---|
| **ERα** | | | | | | | | | | |
| 0 | 4 | 2993 | Cal | 53.21 | 94.34 | 0.94 | 7.2 | 0.943 | 0 | 4.27 |
| | | | Cv | NA | NA | 0.37 | 24.0 | 0.473 | -2.11 | 1.28 |
| 1 | 5 | 1005 | Cal | 88.82 | 98.80 | 0.99 | 3.3 | 0.988 | 0 | 9.29 |
| | | | Cv | NA | NA | 0.71 | 16.3 | 0.665 | 1.85 | 1.89 |
| 2 | 3 | 596 | Cal | 74.99 | 96.87 | 0.97 | 5.3 | 0.969 | 0 | 5.74 |
| | | | Cv | NA | NA | 0.83 | 12.4 | 0.819 | 1.02 | 2.47 |
| 3 | 3 | 419 | Cal | 77.93 | 96.70 | 0.97 | 5.5 | 0.967 | 0 | 5.59 |
| | | | Cv | NA | NA | 0.87 | 10.7 | 0.872 | 0.23 | 2.86 |
| 4 | 3 | 368 | Cal | 75.18 | 96.63 | 0.97 | 5.5 | 0.966 | 0 | 5.54 |
| | | | Cv | NA | NA | 0.89 | 10.2 | 0.923 | -0.09 | 2.99 |
| 5 | 3 | 349 | Cal | 74.58 | 96.52 | 0.97 | 5.6 | 0.965 | 0 | 5.44 |
| | | | Cv | NA | NA | 0.89 | 10.2 | 0.922 | 0.06 | 3 |
| 6 | 3 | 336 | Cal | 74.38 | 96.74 | 0.97 | 5.4 | 0.967 | 0 | 5.62 |
| | | | Cv | NA | NA | 0.90 | 9.7 | 0.915 | 0.19 | 3.16 |
| 7 | 3 | 333 | Cal | 73.56 | 96.83 | 0.97 | 5.4 | 0.968 | 0 | 5.7 |
| | | | Cv | NA | NA | 0.90 | 9.6 | 0.913 | 0.24 | 3.18 |
| **8** | **3** | **332** | **Cal** | **73.61** | **96.81** | **0.97** | **5.4** | **0.968** | **0** | **5.69** |
| | | | **Cv** | **NA** | **NA** | **0.90** | **9.7** | **0.914** | **0.23** | **3.15** |
| **Anti-AR** | | | | | | | | | | |
| 0 | 1 | 2993 | Cal | 24.39 | 72.22 | 0.72 | 13.2 | 0.722 | 0 | 1.93 |
| | | | Cv | NA | NA | 0.55 | 16.8 | 0.584 | -0.62 | 1.51 |
| 1 | 1 | 1043 | Cal | 87.16 | 68.63 | 0.69 | 14.0 | 0.686 | 0 | 1.81 |
| | | | Cv | NA | NA | 0.62 | 15.4 | 0.659 | -0.28 | 1.65 |
| 2 | 4 | 751 | Cal | 90.90 | 97.62 | 0.98 | 3.9 | 0.976 | 0 | 6.58 |
| | | | Cv | NA | NA | 0.66 | 14.6 | 0.77 | 0.27 | 1.74 |
| 3 | 5 | 678 | Cal | 92.72 | 98.91 | 0.99 | 2.6 | 0.989 | 0 | 9.73 |
| | | | Cv | NA | NA | 0.73 | 13.0 | 0.795 | 0.74 | 1.96 |
| 4 | 5 | 662 | Cal | 92.38 | 98.97 | 0.99 | 2.5 | 0.99 | 0 | 9.99 |
| | | | Cv | NA | NA | 0.74 | 12.9 | 0.8 | 0.76 | 1.98 |
| **5** | **5** | **661** | **Cal** | **92.22** | **98.98** | **0.99** | **2.5** | **0.99** | **0** | **10.05** |
| | | | **Cv** | **NA** | **NA** | **0.74** | **12.9** | **0.8** | **0.77** | **1.98** |

Table S12. Performance of the PLS regression of chemical features (abundance) and receptor activity (normalized $EC_{20/50}$ values) detected in PE, PET, PP and PS FCAs and comparison of cross-validated (CV) root mean squared errors (RMSE) and CV determination coefficient ($R^2$) of the initial model 0 and optimized model (VIP filtered). Number of features included in the optimized model and corresponding values for the validation based on 500 iterations with randomly chosen variables. Model 0 included 2993 features.

| Activity | RMSE CV model 0 | RMSE CV optimized model | $R^2$ CV optimized model | Nr features optimized model | PLSR No | RMSE CV random | | $R^2$ CV random | |
|---|---|---|---|---|---|---|---|---|---|
| | | | | | | min | max | min | max |
| PXR | 28 | 22 | 0.64 | 1533 (51%) | 7 | 30 | 31 | 0.26 | 0.32 |
| PPARγ | 21 | 17 | 0.72 | 729 (24%) | 6 | 20 | 22 | 0.53 | 0.61 |
| ERα | 24 | 10 | 0.9 | 332 (11%) | 9 | 29 | 37 | -0.54 | 0.1 |
| Anti-AR | 17 | 13 | 0.74 | 661 (22%) | 6 | 16 | 18 | 0.5 | 0.6 |

Table S13. $AC_{50}$ values of tentatively identified compounds in the optimized PLS regression models co-correlating with the receptor activity.

| PLS model | Compound Name | CAS | $AC_{50}$ | Plastic related | Use |
|---|---|---|---|---|---|
| PXR | triethylene glycol | 112-27-6 | **0.1** | yes | plasticizer, solvent, fragrance |
| | piperine | 94-62-2 | **1.3** | | food additive |
| PPARγ | tetradecanoic acid | 544-63-8 | 118.2 | yes | surfactant |
| | laurocapram | 59227-89-3 | 7.9 | | cosmetics |
| ERα | triphenyl phosphate | 115-86-6 | 7.31 | yes | plasticizer, flame retardant |
| AR | 1-dodecyl-2-pyrrolidinone | 2687-96-9 | 81.5 | tentative | surfactant |
